# In-situ energy budget of needle-leaves reveals shift from evaporative to ‘air cooling’ under drought

**DOI:** 10.1101/2022.04.29.489999

**Authors:** Jonathan D. Muller, Eyal Rotenberg, Fyodor Tatarinov, Itay Oz, Dan Yakir

## Abstract

- The modulation of the leaf energy budget and the balance between its sensible heat (*H*) and latent heat (*LE*) fluxes is vital for vegetation functioning and survival, as it is linked to maintaining leaf temperature below the thermal threshold, an increasingly important mechanism under a drying and warming climate, when evaporative cooling is suppressed.
- Combining measurements and theoretical estimates using a new methodology, we obtained rare and comprehensive energy budgets of leaves on twigs under field conditions in droughted and non-droughted plots of a semi-arid pine forest with low and high evapotranspiration rates, respectively.
- An examination of all components of the needle-leaf energy budget indicated that under the same radiative load, leaf cooling shifts from nearly equal contributions to H and LE in non-droughted trees to almost exclusively H in droughted ones while maintaining a similar leaf temperature.
- This LE-to-H shift in leaves of droughted trees highlights the efficiency of the ‘air cooling’ mechanism in maintaining temperature, which can enhance the resilience of trees to drying conditions. Additionally, leaf energy budgets are a fundamental tool to help understand leaf cooling and aerodynamic resistance under field conditions, and to improve modelling of ecosystem activity and its effect on the climate system.

## 1 Introduction

Leaf energy management, and therefore temperature, affects most physiological aspects of plant and leaf function (Berry and Bjorkman, 1980; Körner, 2006), such as photosynthesis and water loss (O’sullivan et al., 2017; Smith et al., 2019; Richardson et al., 2021; Long et al., 1994; Werner et al., 2002). Indeed, leaf temperature should increase with decreasing evaporative cooling due to a shift towards heat dissipation through sensible heat (*H*), leading to overheating and leaf senescence (e.g., Blonder and Michaletz, 2018; Aparecido et al., 2020; Ruehr et al., 2016) when temperature exceeds the critical limit of biochemical processes (41.5 to 50.8 °C; O’sullivan et al., 2017; Lancaster and Humphreys, 2020; Slot et al., 2021). To avoid this, it is theorised that leaves can sometimes increase or at least maintain stomatal conductance at a high level (Blonder and Michaletz, 2018; Drake et al., 2018). However, this can lead to embolisms due to a water loss through transpiration beyond the plant’s ability to replenish it (Lens et al., 2013; McDowell et al., 2008; McDowell, 2011). In colder climates, processes such as leaf unfolding and frost damage (Bigler and Bugmann, 2018) or the photosynthetic rate are functions of leaf temperature. This importance of temperature has led to increased recent interest in leaf thermal regulation and energy budget modulation (Still et al., 2021; Blonder et al., 2020; Muller et al., 2021a). Our previous results show that the shift to ‘air cooling’ through *H* can be achieved without significantly increasing leaf temperature (Muller et al., 2021a). Indeed, plants modulate the energy budget of leaves through the adaptations to local conditions (e.g. leaf size, shape and orientation, radiation uptake through pigment density) in order to optimise leaf temperature and avoid heat damage (Leigh et al., 2017; Smith, 1978), but how leaves modulate the energy budget to achieve this under changing conditions is currently unknown.

At equilibrium, the energy gained from incoming solar shortwave and thermal longwave radiation is balanced by losses due to thermal radiation, latent heat and sensible heat exchange fluxes, with minor contributions to processes such as biochemical reactions (Gutschick, 2016; Schymanski and Or, 2016), heat storage (except in thick leaves; Schymanski et al., 2013). Plants with a high water availability are efficiently cooled through latent heat exchange (*LE*), while drought is expected to lead to a large increase in H, resulting from a complex interplay between the leaf-to-air temperature difference (Δ*T*_*L−A*_) (Fuchs, 1990) and the aerodynamic resistance to heat exchange (*r*_*H*_). A low *r*_*H*_ is a function of small leaf dimensions and wind speed (Monteith and Unsworth, 2013; Schymanski and Or, 2017) and allows droughted leaves to maintain a low temperature that isn’t significantly higher than with evaporative cooling through *LE* (Muller et al., 2021a).

The leaf energy budget variables can be measured accurately under controlled conditions employing specific radiation and wind regimes (Gutschick, 2016; Schymanski and Or, 2016; Michaletz and Johnson, 2006; Prashar and Jones, 2016): Pioneering work has been carried out at the leaf scale since the 1960s under lab conditions on artificial leaves (Gates, 1962; Gates et al., 1968; Grant, 1984; Schymanski and Or, 2017). Furthermore, the effect of convective heat exchange that affects the sensible heat flux has been studied in depth in wind tunnels (Michaletz and Johnson, 2006; Schuepp, 1993; Gates et al., 1965b; Tibbals et al., 1964). However, leaf-scale energy budget measurements have been difficult to achieve under field conditions. While some have done field studies using devices with small leaf chambers (e.g. LI-COR LI-6400; Sperlich et al., 2019; Schymanski and Or, 2016, these devices do not measure the sensible heat flux and outgoing solar and thermal radiation, and conditions inside the chamber are also not identical to those in free air. Additionally, the effect of neighbouring leaves, either on the same twig or other branches, can’t be taken into account but may crucially change energy budget parameters such as radiation or airflow patterns. To our knowledge, it has therefore not been possible to measure H at the leaf and twig scale under field conditions. Consequently, most studies have focused their efforts on the ecosystem scale, but an in-depth assessment at the leaf- and twig-scale is a crucial gap.

**Aims:** In this study, we used our previously described systems (Muller et al., 2021b,a) to measure the components of the energy budget of needle-leaves on a twig under field conditions in a *Pinus halepensis* forest at the edge of the Negev desert in Israel, during the peak of summer drought. This was done in a manipulation experiment in both a drought-exposed and a summer-irrigated plot. Our objectives were:(a) to examine the leaf energy budget underlying the previously identified paradox of similar incident short- and longwave radiation and leaf-to-air temperature difference in spite of a 20*×* larger LE in irrigated trees, compared with droughted trees (Muller et al., 2021a); (b) to understand the leaf- and branch-scale sensible heat flux at the basis of the observed large ecosystem-scale H flux previously termed the ‘canopy convector effect’ (Rotenberg and Yakir, 2011). Our hypothesis was that the efficient non-evaporative cooling mechanism that keeps needle-leaves of drought-exposed trees similarly cool as highly evaporating ones under the same radiation regime is based on a shift from LE towards ‘air cooling’ using *H*.

### 1.1 Theory of leaf energy budget

The energy budget of leaves on a twig is determined by radiative and non-radiative energy fluxes (mass exchange) between all sides of leaves (i.e., the entire leaf surface) and their surroundings (e.g., Schymanski et al., 2013; Schymanski and Or, 2017; Forseth and Norman, 1993; Ehleringer, 2000; Michaletz et al., 2015). Below, the term ‘leaf’ will refer to multiple needle-leaves on a pine twig, taking the sheltering effect of neighbouring needles into account. The net radiative flux (*R*_*n*_) absorbed by leaves from all directions includes the net absorbed shortwave *S*_*abs*_ (0 to 4000 nm) and net longwave radiation *L*_*n*_ (4 to 100 μm; Ross, 2012) minus fluorescence (*Fl*), while the non-radiative fluxes consist of the sensible (*H*) and latent heat fluxes (*LE*), heat storage (*G*) and energy used in biochemical reactions (*B*), as well as the energy exchanged through the temperature difference between the leaf and sap entering it from the branch (*F*_*S*_; Fig. 1). All variables are defined per total leaf surface area and are summarised in Table 1.

**Table 1.** Symbols used in this study. Values are per two-sided/curved leaf area unless stated otherwise, and temperatures are given in °C unless a different unit is mentioned. Short- (*S*) and longwave radiation (*L*) are in the 0 to 4000 nm and 4 to 50 μm range, respectively.

| Symbol | Description | Unit |
| --- | --- | --- |
| $\alpha$ | Absorptance | — |
| $\alpha_{s,\lambda}$ | Absorptance spectrum (shortwave range) | — |
| $\varepsilon_{d,L}$ | Directional emissivity of a leaf | — |
| $\varepsilon_L$ | Hemispherical emissivity of a leaf | — |
| $\lambda$ | Wavelength | nm |
| $\lambda_w$ | Enthalpy of vaporisation of water | $\text{J mol}^{-1}$ |
| $\rho$ | Reflectance | — |
| $\rho_A$ | Density of air | $\text{g m}^{-3}$ |
| $\rho_L$ | Reflectance (longwave range) | — |
| $\rho_{s,\lambda}$ | Reflectance spectrum (shortwave range) | — |
| $\sigma$ | Stefan-Boltzman constant | $5.670\,374\,419 \times 10^{-8} \text{ W m}^{-2} \text{ K}^{-4}$ |
| $\tau$ | Transmittance | — |
| $\tau_{s,\lambda}$ | Transmittance spectrum (shortwave range) | — |
| $\Delta T_{L-A}$ | Leaf-to-air temperature difference | °C or K |
| $B$ | Energy used in biochemistry | $\text{W m}^{-2}$ |
| $c_{p,C}$ | Heat capacity of carbon | at 20 °C: $0.71 \text{ J g}^{-1} \text{ K}^{-1}$ |
| $c_{p,L}$ | Heat capacity of a leaf | $\text{J g}^{-1} \text{ K}^{-1}$ |
| $c_{p,w}$ | Heat capacity of water | at 20 °C: $4.182 \text{ J g}^{-1} \text{ K}^{-1}$ |
| $C_{w,A}$ | Water concentration in the air | $\text{mol}_{\text{H}_2\text{O}} \text{ m}_{\text{air}}^{-3}$ |
| $C_{w,L}$ | Water concentration in the leaf (i.e. inter-stomatal air space) | $\text{mol}_{\text{H}_2\text{O}} \text{ m}_{\text{air}}^{-3}$ |
| $dT/dt$ | Change in temperature of a leaf over time | $\text{K s}^{-1}$ |
| $d, e$ | Slope and intercept for the empirical calculation of $S_{\uparrow,bL}$ from $S_{\downarrow,aL}$ | — |
| $F_S$ | Heat loss or gain from sap flow into leaves | $\text{W m}^{-2}$ |
| $G$ | Heat storage flux (here: for leaves) | $\text{W m}^{-2}$ |
| $H$ | Sensible heat flux | $\text{W m}^{-2}$ |
| $L$ | Longwave thermal radiant flux (hemispherical) | $\text{W m}^{-2}$ |
| $L_{\downarrow,ac}$ | Downwelling $L$ above the canopy | $\text{W m}^{-2}$ |
| $L_{abs}$ | $L$ absorbed by a leaf | $\text{W m}^{-2}$ |
| $L_{emitted}$ | $L$ emitted by a leaf | $\text{W m}^{-2}$ |
| $L_{in}$ | Incoming $L$ at the leaf | $\text{W m}^{-2}$ |
| $L_n$ | Net $L$ at the leaf | $\text{W m}^{-2}$ |
| $M_w$ | Molar weight of water | $18.015\,28 \text{ g mol}^{-1}$ |
| $m_C$ | Mass of carbon in a leaf | g |
| $m_w$ | Mass of water in a leaf | g |
| $LA$ | Leaf surface area | $\text{m}^2$ |
| $LE$ | Latent heat flux | $\text{W m}^{-2}$ |
| $P$ | Air pressure | hPa |
| $PAR$ | Hemispherical photosynthetically active radiation | $\mu\text{mol m}^{-2} \text{ s}^{-1}$ |
| $R_n$ | Net radiation | $\text{W m}^{-2}$ |
| $r_H$ | Aerodynamic resistance to heat transfer | $\text{s m}^{-1}$ |
| $r_w$ | Resistance to water vapour transfer | $\text{s m}^{-1}$ |
| $S$ | Hemispherical shortwave radiant flux | $\text{W m}^{-2}$ |
| $S_\lambda$ | Hemispherical spectral radiant flux (shortwave) | $\text{W m}^{-2} \text{ nm}^{-1}$ |
| $S_{\lambda,\downarrow,aL}$ | Downwelling $S_\lambda$ reaching a leaf from above | $\text{W m}^{-2} \text{ nm}^{-1}$ |
| $S_{\lambda,\downarrow,ac}$ | Above-canopy downwelling $S_\lambda$ | $\text{W m}^{-2} \text{ nm}^{-1}$ |
| $S_{\lambda,\uparrow,bL}$ | Upwelling $S_\lambda$ reaching a leaf from below | $\text{W m}^{-2} \text{ nm}^{-1}$ |
| $S_{\lambda,\uparrow,in}$ | Mean $S_\lambda$ reaching a leaf from above and below | $\text{W m}^{-2} \text{ nm}^{-1}$ |
| $S_{\lambda,\downarrow,mod}$ | Modelled downwelling $S_\lambda$ | $\text{W m}^{-2} \text{ nm}^{-1}$ |
| $S_{\lambda,\uparrow,mod}$ | Modelled upwelling $S_\lambda$ | $\text{W m}^{-2} \text{ nm}^{-1}$ |
| $S_{abs}$ | Shortwave radiation absorbed by a leaf | $\text{W m}^{-2}$ |
| $S_{abs,upper}$ | Absorbed $S$ by the upper half of a leaf | $\text{W m}^{-2}$ |
| $S_{abs,lower}$ | Absorbed $S$ by the lower half of a leaf | $\text{W m}^{-2}$ |
| $S_{in}$ | Total incoming $S$ at the leaf (mean from above and below) | $\text{W m}^{-2}$ |
| $S_{\downarrow,ac}$ | Downwelling $S$ above the canopy | $\text{W m}^{-2}$ |
| $S_{\downarrow,bc}$ | Downwelling $S$ below the canopy | $\text{W m}^{-2}$ |
| $S_{\uparrow,ac}$ | Upwelling $S$ above the canopy | $\text{W m}^{-2}$ |
| $S_{\uparrow,bc}$ | Upwelling $S$ below the canopy | $\text{W m}^{-2}$ |
| $T_A$ | Air temperature | $^{\circ}\text{C}$ or $\text{K}$ |
| $T_L$ | Leaf temperature | $^{\circ}\text{C}$ or $\text{K}$ |
| $TR$ | Transpiration flux | $\text{mmol m}^{-2} \text{s}^{-1}$ |

**Fig. 1.**
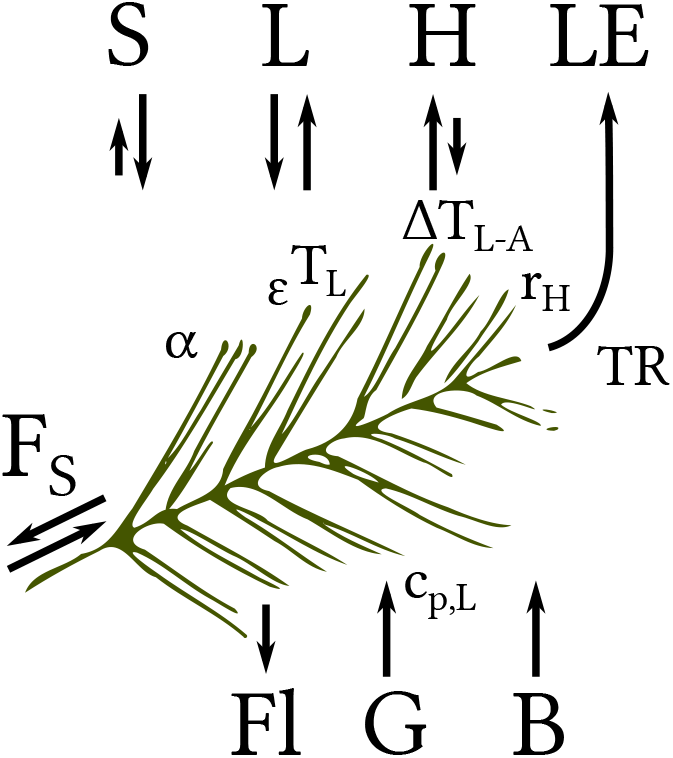
Energy budget of needle-leaves on a twig with the net shortwave (*S*) and net longwave radiation fluxes (*L*), sensible heat (*H*), latent heat (*LE*), heat storage (*G*), energy used in biochemical reactions (*B*), energy emitted through fluorescence (Fl) and energy gained/lost through differences in sap and leaf temperature (*F*_*S*_). S is most strongly affected by the absorptance of leaves (*α*) and their orientation, *L* by their emissivity (*ε*) and temperature (*T*_*L*_), *H* by the leaf-to-air temperature difference (Δ*T*_*L−A*_) and aerodynamic resistance (*r*_*H*_), *LE* by the transpiration (*TR*) and *G* by the leaf heat capacity (*c*_*p,L*_).

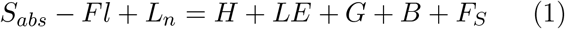

Leaves are required to modulate the terms above to the degree that is possible to maximise assimilation while minimising water loss and avoiding overheating (Fig. 1). This can be achieved through either adaptations in leaf shape and orientation affecting *S*_*abs*_ through mutual leaf shading, as well as the leaf boundary layer which lowers *r*_*H*_ and *T*_*L*_ (Blonder and Michaletz, 2018), or an increased stomatal opening and in turn *TR* which lowers *T*_*L*_ at the cost of a higher water loss. Note that the leaf distribution on a twig plays an equally important role for the boundary layer.

### Net absorbed shortwave radiation (*S*_*abs*_) and spectral effects

Leaves can conceptually be divided into an upper and a lower half to analyse the incident shortwave radiation. Hence, *S*_*abs*_ corresponds to the sum of the direct and diffuse hemispherical *S*_↓,*aL*_ (i.e., from the sky through the canopy) absorbed by the upper half of a total leaf surface (*S*_*abs,upper*_) and the diffuse *S*_↑,*bL*_ reflected from the canopy and ground, absorbed by the lower half of a leaf’s surface (*S*_*abs,lower*_; Eq. 3). In a partly illuminated leaf, the energy budget of the sunlit and shaded parts would theoretically have to be calculated separately, with an additional term for heat conduction between the two parts (Schymanski and Or, 2017). However, this term is neglected here due to the small size of needle-leaves. Measurements of *S* and *L* fluxes are typically per m^2^ of projected surface area, but need to be adjusted to the full leaf surface area (see Methods S1) to correspond to the other heat fluxes.

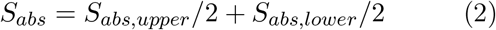

As the absorption of light depends on the wavelength (Jones et al., 2003; Gates et al., 1965a), both the absorptance spectrum of leaves (*α*_*s,λ*_) and the hemispherical shortwave spectral radiant flux reaching leaves (*S*_*λ*_) have to be known or estimated. *α*_*s,λ*_ corresponds to the non-reflected (*ρ*_*s,λ*_) or transmitted (*τ*_*s,λ*_) proportion, i.e. *α*_*s,λ*_ = 1 *− ρ*_*s,λ*_ *− τ*_*s,λ*_. While *ρ*_*s,λ*_ can typically be directly measured using spectrometers, *τ*_*s,λ*_ is difficult to measure, especially for small needle-leaves, and is best obtained from literature databases such as LOPEX93 (Hosgood et al., 1995). The *S*_*λ*_ spectrum is affected by canopy interception and ground reflectance (Fig. 2) and is typically measured using spectrometers or estimated using models. When direct spectral measurements are not available, as in our case, this approach provides a reasonable alternative to realistically adjust the measurements of light intensity of S to the change in spectrum related to solar angle and reflectance from ecosystem elements (e.g. canopy, ground). *S*_*abs,upper*_ and *S*_*abs,lower*_ each corresponds to the integrated hemispherical absorbed *S*_*λ*_ reaching a leaf from above and below (*S*_*λ*,↓,*aL*_ and *S*_*λ*,↑,*bL*_), respectively:

**Fig. 2.**
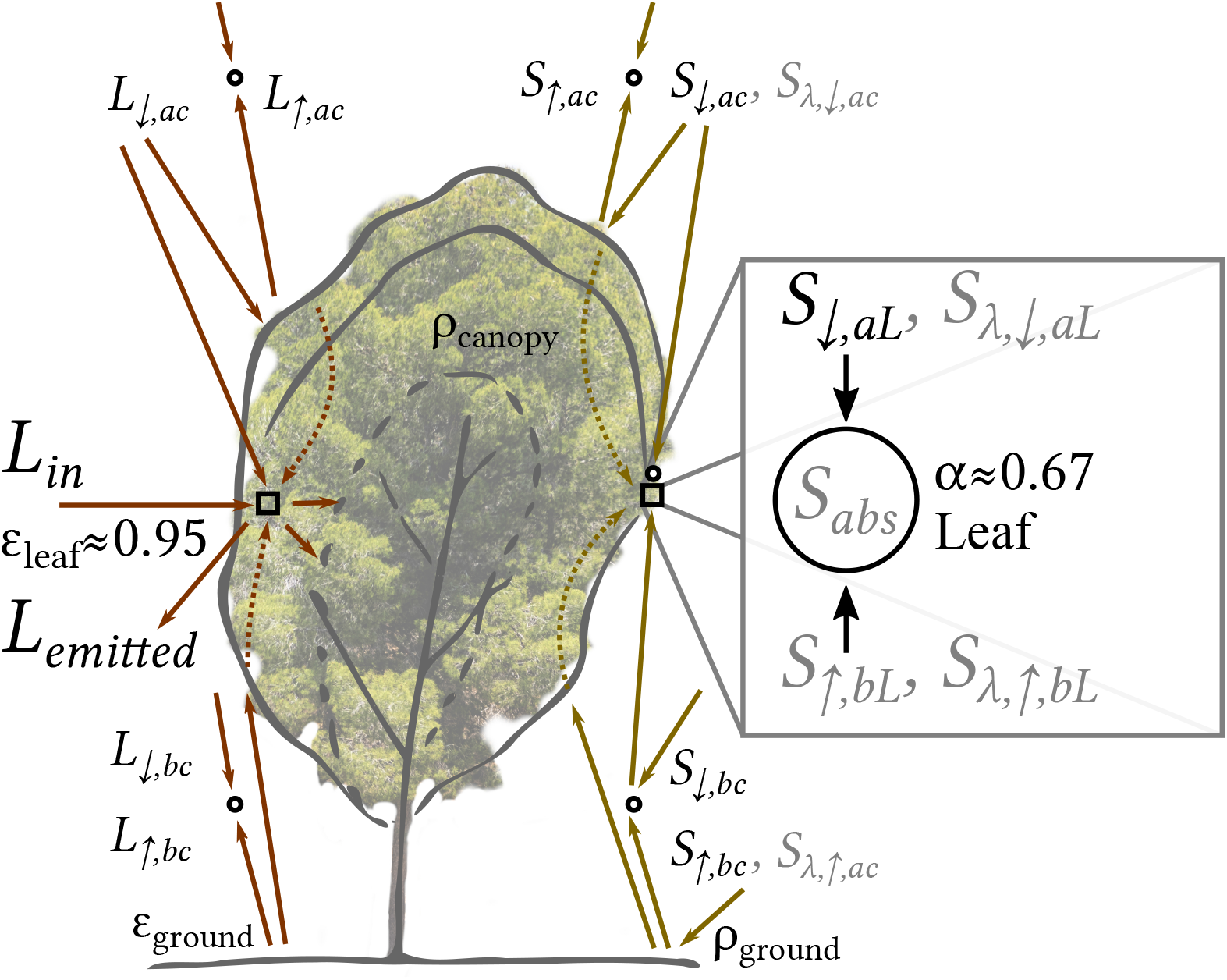
Long- (*L*) and shortwave (*S*) radiation with its spectrum (*S*_*λ*_) reaching a leaf, from above (*ac*), below (*bc*) and within the canopy, with up- (↑) and downwelling (↓) denoting the direction. Measured variables are written in black, while required and calculated ones are in grey. Full-range sensors were available above and below the canopy (locations marked with small circles), but not always near leaves within the canopy. Note that *S* and *L* were measured on the same side of the tree but shown here on different sides for convenience. Finally, *S*_*abs*_ was calculated from the horizontal hemispheres from above and below a leaf, while *L*_*in*_ and *L*_*emitted*_ were measured from a vertical hemisphere integrating values from above and below.

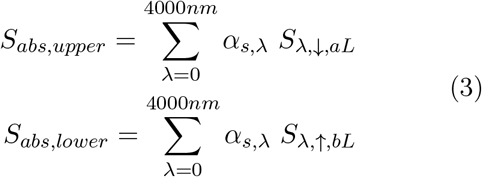

### Net longwave radiation (*L*)

The absorbed *L*_*n*_ corresponds to the incident *L*_*in*_ at the leaf from the environment (i.e. the background longwave radiation), absorbed according to the leaf’s hemispherical absorptivity (i.e. *L*_*abs*_ = (1 − *ρ*_*L*_) *L*_*in*_ = *ε*_*L*_ *L*_*in*_) minus the emitted thermal radiation (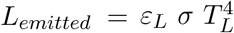; Fig. 2), which is a function of *T*_*L*_:

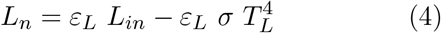

*L*_*in*_, *T*_*L*_ and *ε*_*L*_ have to be accurately measured. Note that the longwave radiation spectrum could also be affected by canopy interception and ground reflectance (similarly to *S*_*λ*_), but it is typically difficult to obtain spectral measurements for this range.

Finally, the combination of *S*_*abs*_ and *L*_*n*_ provides an accurate estimate of *R*_*n*_.

### Sensible heat flux (*H*)

The exchange of sensible (*H*) between the leaves and free air is controlled by convective transport, which is expressed as a function of Δ*T*_*L−A*_ and *r*_*H*_, as well as the air density (*ρ*_*A*_) and heat capacity (*c*_*p,A*_):

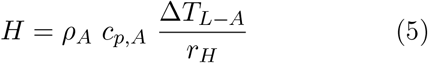

Directly measuring *H* for leaves on a twig is currently impossible under field conditions. However, all the variables of Eq. 5 except for *r*_*H*_ can be measured directly due to the uncertainties related to *r*_*H*_, especially under field conditions when leaves are affected by their neighbours. Previously, *r*_*H*_ has sometimes been calculated from convection measurement in wind tunnel under controlled lab conditions (Tibbals et al., 1964; Gates et al., 1965b; Michaletz and Johnson, 2006). Under field conditions however, *H* can be calculated from the energy budget (Eq. 1). Then, *r*_*H*_ can be obtained by combining Eqs. (1) and (5) (Liu et al., 2007). Note that this dependence on *r*_*H*_ means that ‘air cooling’ through *H* is affected by leaf orientation, geometry, surface properties (e.g. hairs or smoothness) and air flow around leaves and Δ*T*_*L−A*_ (Monteith and Unsworth, 2013).

### Latent heat flux (*LE*)

Measurements of transpiration (*TR*) can be converted to *LE* by simply multiplying it with the enthalpy of vaporisation (*λ*_*w*_, obtained empirically, see Methods S2):

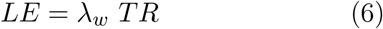

Alternatively, latent heat can also be described similarly to the sensible heat flux, as the difference in water concentration between the leaf 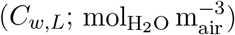 and the air (*C*_*w,A*_) divided by the resistance to water vapour transfer (*r*_*w*_, see Methods S2 & S3).

### Energy used in biochemical reactions (*B*)

A cascading series of partial efficiencies of steps in the photosynthetic pathways leads to losses of the solar radiation reaching leaves (Zhu et al., 2008), resulting in a maximum photosynthetic efficiency of 3.5 % of the incident *S*_*in*_ from all directions under the optimal conditions of times of rapid growth (Blankenship et al., 2011; Zhu et al., 2008). Thus, *B* at a time *t* (*B*_*t*_) is related to that under optimal conditions (*B*_*opt*_) and its net assimilation rate (*A*_*net,opt*_), as well as the current assimilation rate *A*_*net*_ at *t* (*A*_*net,t*_):

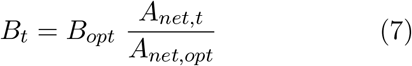

*B*_*opt*_ can be calculated from *S*_*in*_ under optimal conditions (*S*_*in,opt*_):

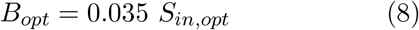

### Heat storage flux (*G*) and flux from sap flow into leaves (*F*_*S*_)

*G* is a function of the change in *T*_*L*_ over time (i.e., *dT*_*L*_*/dt*) resulting from heat transferred into the mass of a leaf (*m*_*L*_). Steady-state leaf temperature is usually reached within 5 min even for 1 mm thick leaves (Schymanski et al., 2013), i.e. leaf heat capacity doesn’t play a role for leaf temperature at longer time scales for needle-leaves (Methods S4). *F*_*S*_ is heat gained or lost by leaves through the mass (*m*_*w*_) of warmer or cooler sap water (*T*_*L*_ *− T*_*S*_) entering leaves due to transpiration (*TR*). Given that the amount of water transported into the leaf is on average equal to the amount of water transpired, the amount of heat transported with the water (*c*_*p,w*_ *m*_*w*_ (*T*_*L*_ *− T*_*S*_) *TR*; Methods S5) is negligible compared to the amount of heat converted to latent heat by evaporation (*λ*_*w*_ *TR*).

## 2 Materials & Methods

### 2.1 Site description & meteorological measurements

The Yatir forest research site is located in a 2800 ha ha afforestation of mainly *Pinus halepensis* trees with a height of *ca*. 10 m and a 50 % canopy cover (measured from Lidar data, June 2019) in the dry southern Mediterranean region, at the northern edge of the Negev desert in Israel (31°20^*′*^49^*′′*^N; 35°3^*′*^7^*′′*^E; altitude 600 to 850 m above sea level; cf. Methods S6; Muller et al., 2021a; Qubaja et al., 2020, 2019).

The research site contains an eddy covariance flux tower *ca*. 50 m away from our location operating since 2001, whose above-canopy environmental sensors (air temperature (°C) at 15 m, 3D sonic anemometer at 18.7 m; R3-100, Gill Instruments, Lymington, United Kingdom) were used for auxiliary measurements of meteorological conditions during our consecutive measurement periods on a half-hourly timescale (Methods S6). Above- (15 m) and below-canopy (2.5 m) up- and downwelling short- (0 to 4000 nm, W m^*−*2^; CM21, Kipp Zonen B.V., Delft, The Netherlands) and longwave radiation (4 to 50 μm, W m^*−*2^; Eppley, Newport RI) were measured at the same location. Leaf-scale measurements contained a mast with a thermal infrared camera facing north (7.5 to 13 μm, calibrated to the 4 to 50 μm range; FLIR A320; FLIR Systems, Wilsonville, Oregon, USA) coupled with a highly reflective Infragold®-coated reference plate (Labsphere Inc., North Sutton NH, USA), as described in Muller et al. (2021b). This system was coupled with a small photodiode (400 to 700 nm; G1118, Hamamatsu Photonics K.K., Japan) installed near the leaves, measuring at 1 Hz. The photodiode was calibrated to the full spectral range (0 to 4000 nm) using a full-range shortwave pyranometer (CM21; Kipp Zonen B.V., Delft, The Netherlands) by placing it near the sensors and measuring in changing environmental conditions (Supplement of Muller et al., 2021a). Adjacent to these leaf temperature measurements, a mast with two sonic anemometers (Windmaster Pro, Gill Instruments, Lymington, United Kingdom) at a vertical distance of 1.5 m of each other measured the sensible heat flux for the bottom and top of the canopy layers, respectively.

All measurements were made during the summer drought period of 2019 (Methods S6) at a height of 5 m, i.e. the middle of the canopy, in a plot with drought-exposed conditions (i.e., control) and one with a summer supplemental irrigation treatment maintaining winter soil water availability (Oz, 2021; Preisler, 2019). Midday conditions (10:00-14:00) were similar during the consecutive measurement periods (Details see Methods S6).

### 2.2 Energy budget variables

#### Leaf absorptance spectrum of shortwave radiation

The absorptance spectrum of leaves (*α*_*s,λ*_) required in Eq. 3 was estimated from the transmittance (*τ*_*s,λ*_, normalised to leaf thickness, cf. Methods S7) and reflectance spectra (*ρ*_*s,λ*_) obtained from means of the 460 fresh broadleaf samples (due to the lack of conifer transmittance data) available in the LOPEX93 database (Fig. 3a) measured in an integrated sphere for the range of 350 to 2500 nm (Hosgood et al., 1995). Some *ρ*_*s,λ*_ measurements of needles were made locally (Fig. S1), but *τ*_*s,λ*_ couldn’t be measured locally and hemispherical effects of *ρ*_*s,λ*_ couldn’t be taken into account. Therefore, these measurements were only used to validate the suitability of the LOPEX93 values for the present study (Fig. S1). Note that plants mostly absorb incoming shortwave radiation in the photosynthetically active range (PAR: 400 to 700 nm; Fig. 3a; Hosgood et al., 1995) due to the effect of this range on photosynthesis, which has previously been studied in detail (e.g. Paradiso et al., 2011; Inada, 1976; McCree, 1971).

**Fig. 3.**
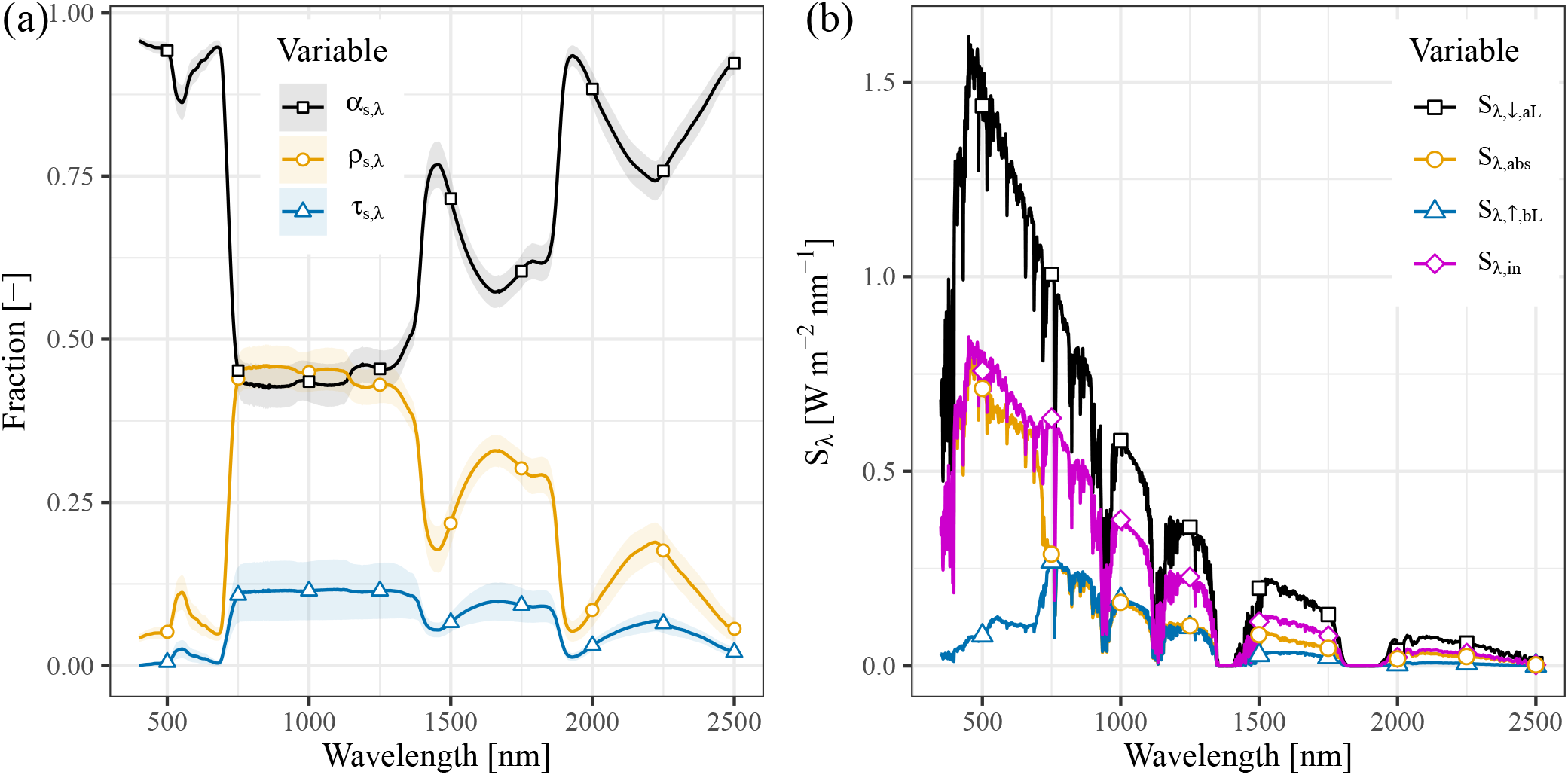
(a) Absorptance (*α*_*s,λ*_, black squares) calculated from reflectance (*ρ*_*s,λ*_, yellow circles) and transmittance (*τ*_*s,λ*_, blue triangles) spectra, averaged from all fresh broadleaf samples in the LOPEX93 database (Hosgood et al., 1995). Note that no transmittance data is available for needle-leaves, hence the usage of broadleaf data. The shaded areas depict standard deviations; (b) Spectrum of shortwave radiation, adjusted from the SMARTS2 model (Gueymard, 1995, cf. Methods S8), reaching the leaf from above (black squares), from below (blue triangles), in total (purple losanges), and absorbed fraction (yellow circles) on an example of midday conditions (2019-07-10 11:18)

#### Up- and downwelling shortwave radiation measurements

Since the spectrum of incoming shortwave radiation from above and below a leaf (*S*_*λ*,↓,*aL*_ and *S*_*λ*,↑,*bL*_) could not be measured directly in our conditions, a number of steps were required to obtain it. The incident light spectrum was estimated using the SMARTS2 clear-sky spectral model (Gueymard, 2005, 1995; Myers and Gueymard, 2004), initialised with environmental data measured above canopy at our Eddy Covariance flux tower, (i.e. latitude, longitude, altitude, date & time, air temperature, relative humidity, atmospheric CO_2_ concentration and ecosystem albedo). The SMARTS2 model was adjusted to above- and within-canopy conditions using integrated measurements (0 to 4000 nm) of above- and below-canopy half-hourly mean hemi-spherical shortwave radiation (*S*_*λ*,↑,*ac*_, *S*_*λ*,↑,*bc*_, *S*_*λ*,↓,*ac*_ and *S*_*λ*,↓,*bc*_) at the EC tower. Measurements of the upper hemisphere of incoming shortwave radiation at the leaf from above (*S*_*λ*,↓,*aL*_) were done with a calibrated PAR photodiode. The detailed calculation method of *S*_*abs*_ is described in Methods S8, and Fig. 3b shows how the spectrum of shortwave radiation is reduced from above the canopy (*S*_*λ*,↓,*aL*_) to the mean from above and below a leaf (*S*_*λ,in*_) and its absorption (*S*_*λ,abs*_).

#### Net longwave radiation (*L*_*n*_) and leaf temperature (*T*_*L*_)

Both *L*_*in*_ (corresponding to back-ground thermal radiation from a south-oriented vertical hemisphere originating from the ground, canopy and sky; Fig. S1 of Muller et al., 2021b) and *T*_*L*_ were obtained using our previously described ‘dual reference’ method (Muller et al., 2021b). Note that this measurement method of *L*_*in*_ supplies the integrated value from above and below leaves.

The directional emissivity of leaves (*ε*_*d,L*_) was measured in the lab for the 4 to 50 μm range at an uncertainty of <0.5 % (Vishnevetsky et al., 2019). However, the total emitted longwave radiation from a surface is a function of the hemispherical emissivity (*ε*_*L*_) because energy is lost to the entire hemisphere. *ε*_*L*_ is lower by 0.026 to 0.035 than *ε*_*d,L*_ in broadleaves (Vishnevetsky et al., 2019). As *ε*_*L*_ could not be measured for pine needles, we subtracted a value of 0.03 from their measured *ε*_*d,L*_, resulting in *ε*_*L*_ ≈ 0.92 and a reflectivity of 1 *− ε*_*L*_.

A set of fine thermocouples installed in a radiation shield (Model 41003; R. M. Young Company, Traverse City MI, USA) *ca*. 2 to 3 m away from the leaves provided *T*_*A*_, while *T*_*L*_ and *L*_*in*_ were both measured using our IR camera-based system (Muller et al., 2021b).

#### Latent heat flux (*LE*)

Twig-scale gas exchange chambers (Oz, 2021; Preisler, 2019) adjacent to the branches used for *T*_*L*_ measurements (Muller et al., 2021a) were used to calculate the evaporative flux of transpiration (*TR*) per unit leaf surface area from 4 twigs, using an advanced Python script (Muller and Oz, 2020). This flux was then converted to *LE* according to Eq. 6 & (S4). Note that the same chambers also provided the CO_2_ flux necessary to estimate assimilation.

#### Energy used in biochemical reactions (*B*)

PAR sensors (SQ-500-ss; Apogee Instruments Inc., Logan UT, USA) were installed in November 2020 next to the twig-scale gas exchange chambers. A 13 % attenuation from the chamber Plexiglas walls was measured during calibration and applied to the PAR data. Then, this PAR data was converted to the full shortwave radiation spectrum using the above-mentioned calibration, thus providing *S*_↓,*aL*_ inside the chambers. The highest 0.5 % of net assimilation during the growing season (January to May 2021) under irrigated conditions was then extracted. This data was used to calculate the energy used in biochemical reactions under optimal conditions (*B*_*opt*_) as 3.5 % of *S*_↓,*aL*_ (Eq. 8). Then, the reduction in assimilation compared to *B*_*opt*_ was calculated from net assimilation values during the measurement period of 2019 (Eq. 7). Note that *B*_*opt*_ is likely overestimated because *S*_↑,*bL*_ could not be considered and because the value of 3.5 % is likely to be too high, even for well-irrigated *P. halepensis*.

#### Fluorescence (*Fl*)

Estimates of *Fl* were modelled using radiation conditions measured at our site (cf. Methods S9). Considering its low magnitude (see Results), *Fl* was ignored in further energy budget calculations.

#### Heat storage flux (*G*) and flux from sap flow into leaves (*F*_*S*_)

Needle-leaf wet and dry weight were measured monthly in irrigated and drought-exposed plots, both in pre-dawn and midday conditions. Due to the small dimensions of needle-leaves (here 0.8 to 1 mm diameter), leaf temperature was assumed to be equivalent to leaf surface temperature. For *F*_*S*_, measurements of branch sap temperature were not available, but a sensitivity analysis was performed, showing that it is negligible (See Methods S5).

#### Sensible heat (*H*)

To our knowledge, there is currently no way to directly measure H at the leaf scale in the field. Hence, the sensible heat flux was calculated as a residual of the energy budget (Eq. 1). Note that *G* was approximated as 0.7 % of *R*_*n*_ (longterm mean), while *Fl* and *F*_*S*_ were ignored as they were considered negligible.

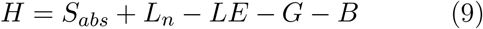

### 2.3 Data processing & filtering

For low *S*_*abs*_, *L*_*n*_ and *LE, H* could not be accurately estimated. Hence, calculated values of *r*_*H*_ (reverse of Eq. 5) are also not valid. This data needs to be identified and excluded from the analysis to avoid errors in the resulting calculations of *H* and *r*_*H*_. For instance, when *LE* ≈ 0 (drought) and *S*_↓,*aL*_ = 0 (night), *H* can only be calculated for *L*_*n*_ > *±*3.45 W m^*−*2^ (i.e., the accuracy of *L* measurements).

Therefore, data was excluded from analysis according to the following criteria: (a) Data with a daily maximum of *S*_↓,*aL*_ < 500 W m^*−*2^; (b) Data with Δ*T*_*L−A*_ < 0.00 *±* 0.28 °C due to the system accuracy (Muller et al., 2021b). (c) Due to the low magnitude of *LE* at night, average night-time values were used for each of the two treatments (drought-exposed and irrigated) for the entire measurement period.

Welch *t* -tests were performed when comparing bins of data to determine the significance of potential differences.

### 2.4 Source of uncertainties in energy budget calculations

Our energy budget data contains a number of uncertainties:(a) The attenuation of shortwave light spectrum due to canopy interception from above (*S*_*λ*,↓,*aL*_) and below the leaf (*S*_*λ*,↑,*bL*_) was estimated, not measured. The spectrum of *S*_*λ*,↓,*aL*_ was estimated from *S*_*λ*,↓,*ac*_ (cf. Methods S3) and that of *S*_*λ*,↑,*bL*_ from the reflected *S*_*λ*,↓,*ac*_ on the ground and leaves. Spectrometers would improve these values in future work. (b) All data of *S*_*abs*_ corresponds to horizontally oriented leaves, even though other leaf orientations could reduce radiation load. Note that for longwave radiation, no effect of leaf orientation is expected. (c) The intensity of *S*_*λ*,↑,*bL*_ was empirically estimated at our EC tower location due to the lack of a down-looking sensor close to the studied leaves, but may not be the same at our measurement location. (d) The SMARTS2 model contains data from 280 to 4000 nm, but the final data used in calculations was equivalent to absorptance spectrum (350 to 2500 nm) available in the LOPEX93 database. This range contained 97 % of the energy of the 280 to 4000 nm range, i.e. *ca*. 3 % uncertainty. (e) Branch chamber walls affect the entire radiation regime (i.e., *S* and *L*), but only the *ca*. 13 % reduction of PAR could be measured, leading to an uncertainty of transpiration measurements estimated at 10 to 15 % for averaged fluxes. Due to its magnitude, this uncertainty was only relevant for *LE* in irrigated trees. (f) *G* and *Fl* were measured at the leaf scale and *T*_*L*_, *S*_*abs*_, *L*_*n*_, *B* and *LE* a twig scale (adjusted to units of leaf surface). Due to the small leaves and low density of conifers in Yatir forest, we assume that the results also represent the twig scale.

## 3 Results

### Radiative energy fluxes

The net twig-scale absorbed radiation (*R*_*n*_) could be provided with a high confidence since it was based on the individually measured short- and longwave radiation fluxes and accounted for spectral effects. Table 2 shows a summary of the radiation fluxes from the top of the canopy to the leaf scale in the middle (5 m) in both treatments during the time of peak *S*_↓,*ac*_ (10:00-12:00 due to the usage of winter time), with comparable total shortwave radiation load between treatments (*S*_*total*_ = (*S*_↓,*aL*_ + *S*_↑,*bL*_)*/*2; Table 2) and similar mean Δ*T*_*L−A*_ in droughted and irrigated trees of 2.76*±*0.70 and 2.79*±*0.69 °C, respectively (*P* = 0.69). Therefore, canopy interception reduced the above-canopy *S*_↓,*ac*_ from about 900 W m^*−*2^ to the downwelling *S*_↓,*aL*_ at the leaf of *ca*. 575 W m^*−*2^, on average (Fig. 4a, Table 2). *S*_↑,*bL*_ was much lower at *ca*. 85 W m^*−*2^ (Table 2) as a result of ground reflectance and canopy interception. Leaf radiation load is further reduced since only 67 % of the total incoming shortwave radiation is absorbed (24 % and 6 % are reflected and transmitted, respectively, with a 3 % uncertainty). This led to a peak of mean daytime *S*_*abs*_ of 220 *±* 24 and 223 *±* 59 W m^*−*2^ (not significantly different, *P* = 0.58) and mean of daily maximum values of 237*±*17 and 257*±*23 W m^*−*2^ in the drought-exposed and irrigated treatments, respectively. Note that *Fl* reached maximum daytime values under the highest incoming solar radiation regime (*ca*. 900 W m^*−*2^) of merely 0.2 W m^*−*2^.

**Table 2.** Summary of radiative fluxes (W m^*−*2^) during peak hours (10:00-12:00) of downwelling above-canopy *S* (*S*_↓,*ac*_), downwelling *S* from above a leaf (*S*_↓,*aL*_), upwelling *S* from below a leaf (*S*_↑,*bL*_), mean incident *S* on a leaf from both sides (*S*_*in*_), absorbed *S* (*S*_*abs*_), incident *L* (*L*_*in*_), *L* absorbed (*L*_*abs*_) and emitted by a leaf (*L*_*emitted*_), net *L* (*L*_*n*_) as well as net radiation (*R*_*n*_ = *S*_*abs*_ + *L*_*abs*_ *− L*_*emitted*_) and the sum of absorbed radiation components (*S*_*abs*_ + *L*_*abs*_). Note that we use UTC+2h (winter time) throughout the year for consistency.

| Variable | Drought-exposed | Irrigated |
| --- | --- | --- |
| $S_{\downarrow,ac}$ | $892 \pm 59$ | $994 \pm 37$ |
| $S_{\downarrow,aL}$ | $573 \pm 62$ | $580 \pm 154$ |
| $S_{\uparrow,bL}$ | $90 \pm 9$ | $91 \pm 24$ |
| $S_{in}$ | $332 \pm 36$ | $335 \pm 89$ |
| $S_{abs}$ | $220 \pm 24$ | $223 \pm 59$ |
| $L_{in}$ | $370 \pm 8$ | $365 \pm 9$ |
| $L_{abs}$ | $350 \pm 8$ | $345 \pm 8$ |
| $L_{emitted}$ | $349 \pm 8$ | $354 \pm 9$ |
| $L_n$ | $1 \pm 2$ | $-8 \pm 3$ |
| $R_n$ | $221 \pm 25$ | $210 \pm 59$ |
| $S_{abs} + L_{abs}$ | $572 \pm 27$ | $565 \pm 59$ |

**Fig. 4.**
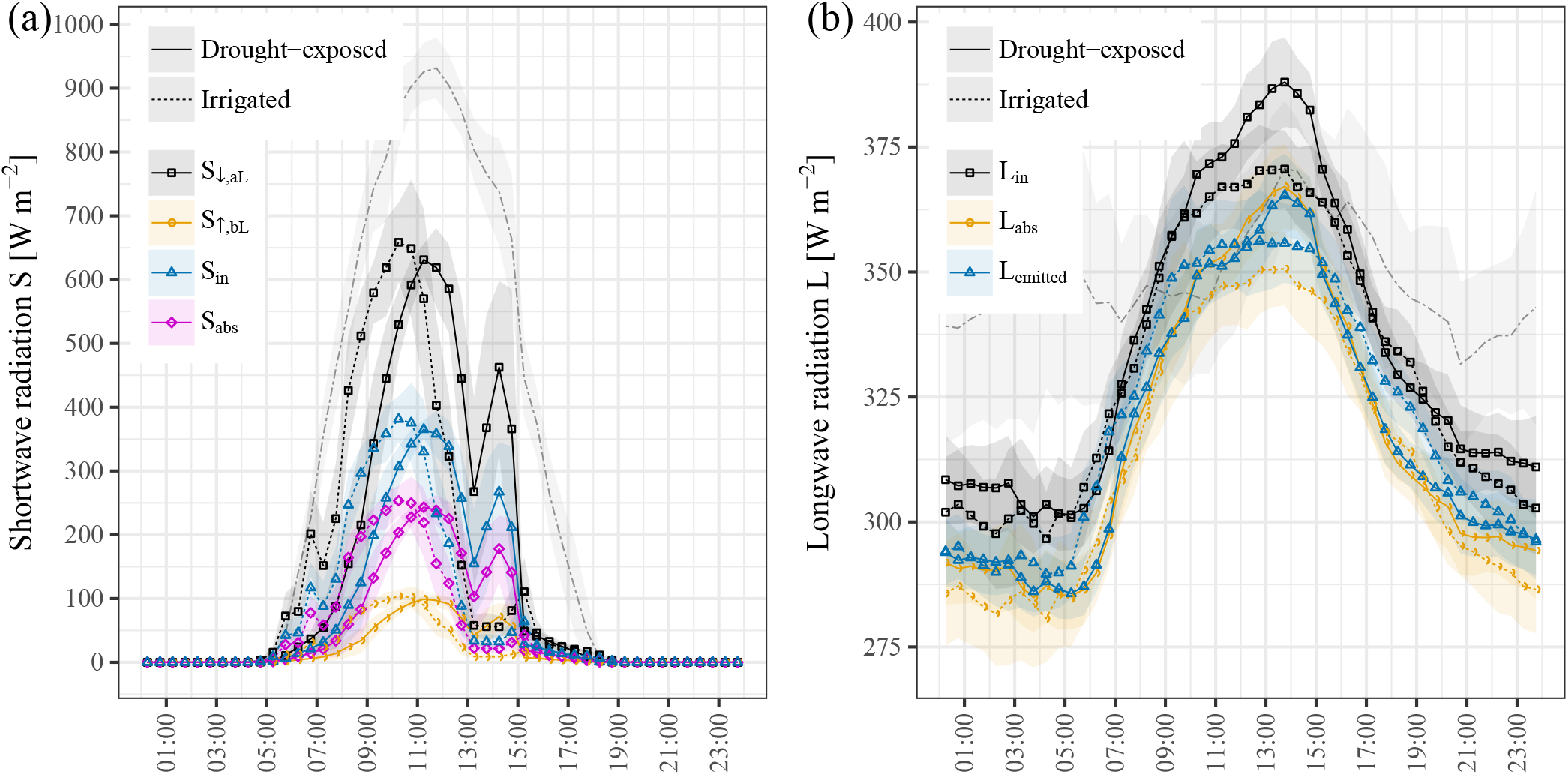
Diurnals of 30 min means of radiation (Aug. & Sept. 2019) in the drought-exposed (solid line) and irrigated treatments (dashed), of (a) shortwave solar radiation: Downwelling above canopy in both treatments (*S*_↓,*ac*_; grey dashed), incident *S* reaching the leaf from above (*S*_↓,*aL*_; black squares) and from below the twig (*S*_↑,*bL*_; yellow circles) and the total incident *S* per total leaf area (i.e., mean of both sides of a leaf; *S*_*in*_; blue triangles), as well as the net absorbed non-reflected *S* (*S*_*abs*_; purple lozenges) and (b) longwave thermal radiation: Downwelling *L* above the canopy in both treatments per flat area (*L*_↓,*ac*_; dashed grey) and, adjusted to the entire curved leaf surface area: incoming from the background (*L*_*in*_; black squares), absorbed (*L*_*abs*_; yellow circles) and emitted by the leaves (*L*_*emitted*_; blue triangles). Shading represents standard deviations. Adjustments to the entire leaf surface area are discussed in Methods S1.

The amount of thermal radiation adjusted to the curved leaf area and absorbed by them (*L*_*abs*_; 95 % of *L*_*in*_) was nearly equal to *L*_*emitted*_ (Table 2, Fig. 4). This was partly due to the measurement location in the middle of the canopy, where a great part of *L*_*in*_ is from the surrounding warm ground and canopy. Therefore, *L*_*n*_ remained near zero in the drought-exposed plot and was on a daily average 0.1 *±* 3.9 W m^*−*2^ (mean of daily max.: 6.6 *±* 3.2 W m^*−*2^, i.e. *L*_*in*_ *> L*_*emitted*_) with only few positive values during the day (*S*_↓,*ac*_ *>* 20 W m^*−*2^). In the irrigated plot, *L*_*n*_ was lower with mean values of *−*9.5 *±* 4.9 W m^*−*2^ (daily max.: 0.0 *±* 7.1 W m^*−*2^; both daily means and mean of maxima: *P <* .001). This significant difference between treatments is most likely caused by the slightly lower soil temperature in the irrigated treatment due to evaporation (see Muller et al., 2021b). Consequently, *R*_*n*_ was almost exclusively determined by *S*_*abs*_ in both treatments, with an approximately *−*10 W m^*−*2^ offset between treatments resulting from negative *L*_*n*_ in the irrigation treatment. Note that *L*_↓,*ac*_ was sometimes larger than *L*_*in*_, *L*_*abs*_ and *L*_*emitted*_ (Fig. 4b) because it corresponded to the incident thermal radiation on a projected surface area, while the latter were adjusted to the curved leaf surface area.

### Full energy budget

At night, *R*_*n*_ was negative in the irrigated treatment (*−* 9 *±*4 W m^*−*2^) due to the negative *L*_*n*_, but remained near zero in drought-exposed trees (*−*3 *±* 3 W m^*−*2^). During the day, *R*_*n*_ reached similar maxima in both treatments (213 *±* 46 and 195 *±* 68 W m^*−*2^; Table 2).

Maximum half-hourly estimated values of *G* (0.06 *±*0.16 W m^*−*2^ or <0.002 % of *R*_*n*_ on average), *B* (5.95 *±*1.82 W m^*−*2^ or <1.15 % and 0.03 *±*0.02 W m^*−*2^ or <0.01 % in irrigated and droughted treatments) and emitted as fluorescence (*Fl*; 0.17*±*0.01 W m^*−*2^ or <0.1 % of *R*_*n*_ in both treatments) were considered negligible and were omitted in further calculations.

*H* was calculated as the residual of the energy budget (Eq. 9). In the drought-exposed treatment, *R*_*n*_ was almost exclusively dissipated as *H* (peak daytime mean: 222*±*25 W m^*−*2^, Fig. 5a). *LE* was indeed very low under drought-exposed conditions (twig-scale: 5.4 ± 4.9 W m^*−*2^, <2.5 % of *R*_*n*_), which is further emphasized by the low ecosystem-scale flux (36 ± 38 W m^*−*2^; Methods S3; Note that the ecosystem scale includes both the considerably larger soil evaporation and transpiration of all leaves, including those in full sun, and the bark evaporation (Lintunen et al., 2021)). In the irrigated treatment, twig-scale *H* was >100 W m^*−*2^ lower during peak daytime activity (129*±*55 W m^*−*2^) since significant energy dissipation occurred through latent heat of evaporation (71.8*±*17.0 W m^*−*2^; Fig. 5b). *H* was near zero at night in the drought-exposed plot but below zero in the irrigated plot, due to the negative *L*_*n*_ and positive *LE* in this treatment. Overall, *H* under irrigation remained at less than ½ of the level of the droughted treatment. Note that the moisture content of the air can affect *H*, but this effect was smaller than *ca*. 2.8 W m^*−*2^ (See Notes S1).

**Fig. 5.**
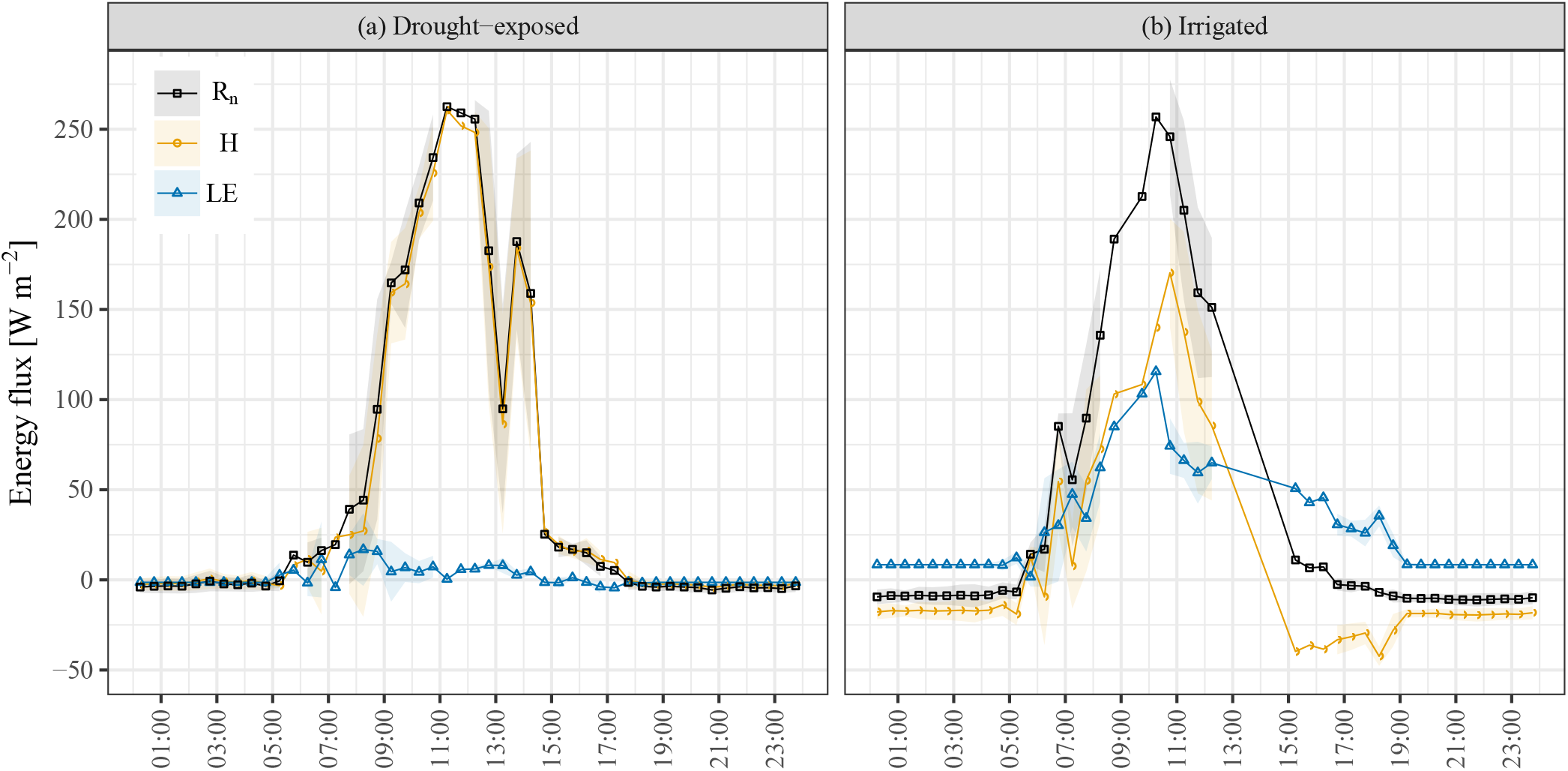
Diurnals of 30 min means of energy fluxes at leaves on a twig of a droughted and irrigated tree during summer 2019, of net absorbed radiation (*R*_*n*_), latent heat (*LE*) and sensible heat (*H*). Heat storage (*G*) and energy converted in biochemical reactions (*B*) are omitted due to their small magnitude.

## 4 Discussion

The detailed partitioning of the leaf energy budget is an important tool for understanding many aspects of its physiological activities (Schymanski et al., 2013). Previously, the leaf energy budget has mostly been measured in lab conditions (Vialet-Chabrand and Lawson, 2019; Schymanski and Or, 2016), through leaf models (Chiariello, 1984; Okajima et al., 2012), or under highly controlled conditions in the field (Knoerr and Gay, 1965; Sperlich et al., 2019). Our setup enabled us to measure and calculate all the components of the energy budget of needle-leaves on twigs in-situ. Our Aleppo pine study site provided the opportunity to carry out these measurements in both drought-exposed trees with nearly zero transpiration, and in trees provided with a summer supplemental irrigation corresponding to winter soil water contents and having a substantial transpiration flux. The energy budgets we report are based on a combination of conventional and novel measurements, complemented by calculated and modelled values, based on the extensive literature available on the topic. This allowed us to account for the spectral dependence of the radiative energy fluxes, determining the emissivity of the relevant surfaces, and obtaining an unprecedented accuracy of estimates of leaf temperature and leaf-to-air temperature differences, shown in our previous publications (Muller et al., 2021a,b). In summary, leaf temperatures remained within *ca*. 3.5 °C of air temperature in both treatments (mean: 2.8 °C), while assimilation rates were up to 7 μmol m^*−*2^ s^*−*1^ in irrigated and near zero in droughted trees (Figure S2).

### Leaf energy budget

Figure 6 provides a summary of all the elements in this budget, as also discussed in more detail in the theory section. It focuses on the main results binned for conditions when the net radiative energy dissipated by nonradiative means (*R*_*n*_) was in the peak range of 200 to 250 W m^*−*2^ (i.e., *S*_↓,*ac*_: 891 *±* 82 W m^*−*2^ and *S*_*in*_: 340*±*24 W m^*−*2^) resulting in a Δ*T*_*L−A*_ without a significant difference. This summary provides a rare characterization of needle-leaf energy budget in our study, including the following main elements: (a) About 30 % and 5 % of the incident shortand longwave radiation were reflected and transmitted by the leaf surface, respectively. (b) The total absorbed radiative energy consisted of *ca*. 40 % shortwave radiation, of which 80 % was in the PAR range (400 to 700 nm, Fig. 3b), and *ca*. 60 % was longwave radiation. Nearly the same amount of longwave radiation as is absorbed is re-emitted, leading to a small *L*_*n*_ (Droughted: 1.7 *±*3.1 W m^*−*2^; Irrigated: *−*10.3 *±*3.3 W m^*−*2^) but a large *S*_*abs*_ (Droughted: 221 *±*15 W m^*−*2^; Irrigated: 235*±*15 W m^*−*2^). This is expected as the measurements were carried out in the middle of the canopy where a great part of *L*_*in*_ is from the surrounding leaves at a similar temperature. While the absorption of longwave radiation is difficult to modify, leaves can reduce the amount of absorbed shortwave radiation through adaptations of their pigmentation, shape and/or orientation (Leigh et al., 2017; Smith and Carter, 1988; Monteith, 1973; Landsberg and Thom, 1971; Michaletz and Johnson, 2006; Gates, 1962). Note also that the magnitude of *L*_*emitted*_ was nearly identical between treatments because leaf temperature and emissivity were similar too at our study site (Muller et al., 2021a). (c) Only a small fraction of the absorbed radiative energy is used in biochemistry (*B <* 1%), or stored (*G*, negligible) in both treatments. To maintain the observed low Δ*T*_*L−A*_ in steady state (Muller et al., 2021a), the rest of the energy must be dissipated radiatively through thermal radiation, or non-radiatively through heat fluxes (sensible and latent). (d) The energy dissipation from a needle-leaf in this semi-arid climate is split into ⅔ emission of longwave radiation and ⅓ through the non-radiative *H* and *LE*. The partitioning of the *H* : *LE* fluxes was 22:17 in irrigated and 37:1 in leaves of droughted trees, highlighting the shift from evaporative to non-evaporative cooling. While *LE* can be dynamically adjusted by plants through stomatal opening, *H* is a parameter that more directly depends on aerodynamic resistance to heat transfer (*r*_*H*_). This parameter, in turn, depends on wind speed, leaf size, geometry and sheltering from neighbouring leaves (Monteith and Unsworth, 2013; Schymanski and Or, 2017; Blonder and Michaletz, 2018), the latter of which can only slowly be adjusted to environmental conditions through growth or leaf senescence.

**Fig. 6.**
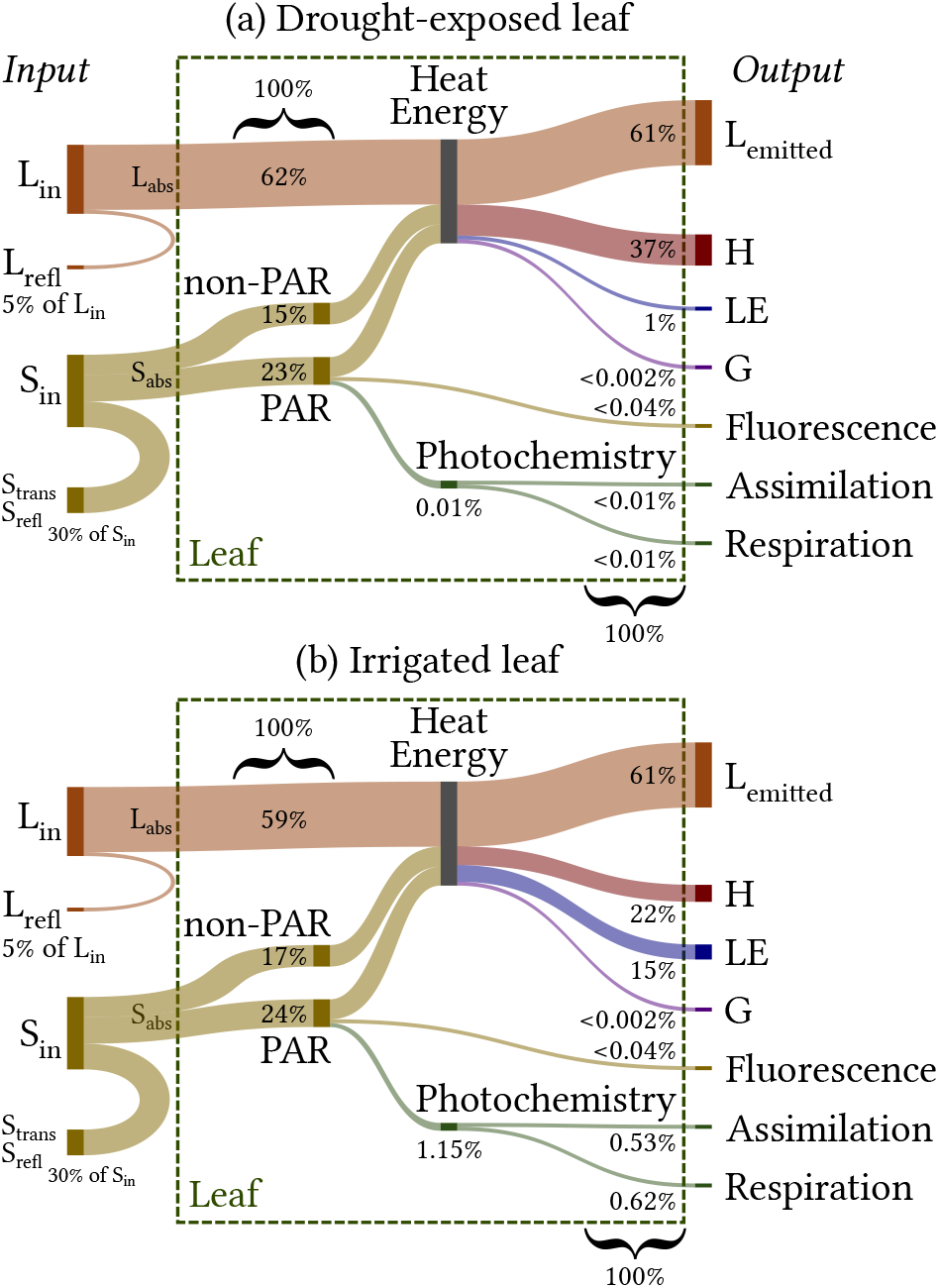
Midday in-situ energy budget of needle-leaves by total surface area, indicated as a proportion of total absorbed radiation (*S*_*abs*_ + *L*_*abs*_), in (a) drought-exposed and (b) experimentally irrigated conditions (576*±*18 and 577 *±*21 W m^*−*2^, respectively): Incoming short- (*S*_*in*_) and longwave radiation (*L*_*in*_) is either reflected (*S*_*refl*_ *& L*_*refl*_), transmitted (*S*_*trans*_) or absorbed. Absorbed shortwave radiation (*S*_*abs*_), is divided into photosynthetically active radiation (PAR) that is used in photochemistry, emitted through fluorescence (*Fl*) and dissipated thermally. Absorbed longwave radiation (*L*_*abs*_) goes to heat entirely. Heat is emitted as longwave radiation (*L*_*emitted*_), released as sensible (*H*) and latent heat (*LE*) and a small amount is stored (*G*) on a half-hourly timescale. All these components are affected by leaf temperature (*T*_*L*_). Note that *R*_*n*_ consists of *S*_*abs*_ + *L*_*abs*_ *− L*_*emitted*_.

Field studies of leaf energy budgets are scarce due to technical challenges of measurements in the field, especially on trees. Therefore, most studies rely on Eddy Covariance for plot-scale measurements. One study on a maize field estimated that 46 % of *R*_*n*_ is allocated to *LE* and 32 % to *H*, whereas the remaining energy is attributed to soil heating and evaporation (Brown and Covey, 1966). While not providing a full energy budget, Blonder et al. (2020) showed in a broad study that several environmental and functional traits can influence the leaf energy budget under field conditions and in different species and ecosystems, but that these are not sufficient to infer leaf energy budgets without detailed work such as advocated for by Monteith and Unsworth (2013) and presented from measurements here. Due to the difficulty of measuring all the leaf energy budget variables, many studies merely discuss the potential energy budget implications of their leaf temperature measurements, often in relation to wind speed (e.g., Leigh et al., 2017; Michaletz et al., 2016; Blonder and Michaletz, 2018; Slot et al., 2021; Gates et al., 1968; Gates, 1962). Finally, studies such as Schymanski et al. (2013); Schymanski and Or (2016) relied on extensive model simulations and showed, for instance, that for 3 cm large leaves at 0.5 m s^*−*1^ wind speed, evaporative cooling would be needed under steady state to prevent heat damage at air temperatures above 34 °C for an irradiance below 600 W m^*−*2^ (Schymanski et al., 2013), while it is required from 300 W m^*−*2^ at air temperatures above 41 °C. Nevertheless, these models simulate variable, and contrasting, partitioning in leaf energy budgets between *H* and *LE* across species. Underpinning these studies, our detailed and relatively unique in-situ twig-scale energy budgets provide a way to quantify the mechanisms underlying the efficient non-evaporative cooling of plant leaves under drought that keeps leaves at similar temperatures as highly evaporating ones and relies on a shift from the latent to the sensible heat flux.

### Aerodynamic resistance (*r*_*H*_)

The detailed energy budget discussed above shows that the main difference between the droughted and irrigated trees was in the suppression of *LE* and the large *H* flux in the droughted trees. *H* is often assumed to reflect surface temperature, or the surface-to-air temperature gradient, but it was shown that these terms were almost identical in the two treatments. Considering that *H* is a function of Δ*T*_*L−A*_ and the resistance to heat transfer *r*_*H*_ (Eq. 5), the large *H* in the droughted trees can only be explained by a reduction in *r*_*H*_ (Note that the similar *T*_*L*_ also rules out an increased radiative energy dissipation). Owing to our ability to provide data on Δ*T*_*L−A*_ and *H*, we could estimate *r*_*H*_ of bunches of leaves on a twig under field conditions. Under similar radiation load during peak activity hours and wind regime (Muller et al., 2021a), leaf-scale *H* of the droughted trees was double that of irrigated ones (Fig. 6), while Δ*T*_*L−A*_ wasn’t significantly different. Hence, we have to assume that *r*_*H*_ is ca. 2*×* smaller (Droughted: 25 *±* 27 and irrigated: 47 *±* 51 s m^*−*1^, *P <* 0.05). How a shift from ca. 50 % evaporative cooling (*ca*. 50 % energy dissipation via *LE*) to ‘air cooling’ (i.e., >96 % energy dissipation via *H*) occurs while Δ*T*_*L−A*_ remains unchanged is an open question. However, the leaf energy budget and resulting ability to calculate *r*_*H*_ is an important tool to improve our understanding of leaf-to-ecosystem energy management.

### Leaf energy budget across the canopy height

Using previously available data and detailed solar and thermal radiation measurements above and below the canopy (Rotenberg and Yakir, 2011) from the Eddy Covariance tower, we were able to provide energy budget estimates for the projected leaf area for the top and bottom of the canopy under drought. The magnitude of the total radiation load and lack of *LE* highlights the necessity of leaves in dry environments to be cooled convectively throughout the canopy through an efficient air flow, as predicted by Defraeye et al. (2013). Leaves at the top of the canopy (*ca*. 10 m) during summer midday absorbed 100 *±* 29 W m^*−*2^ more shortwave radiation than at the bottom (*ca*. 2.5 m), but simultaneously experienced significant radiative cooling (negative *L*_*n*_) of 57 6 W m^*−*2^ (Table 3) as their upper surface is exposed to the cooler atmosphere (*L*_*in*_). This led to a comparable net absorbed radiation load at both heights (*R*_*n*_ ≈ 265 W m^*−*2^) and no significant difference in energy dissipated through *H* throughout the canopy height (*ca*. 265 W m^*−*2^, *P* = 0.39), in spite of significant turbulence differences (shear velocity *u*_*_: top, 0.48 *±* 0.16 m s^*−*1^; bottom, 0.23 *±* 0.04 m s^*−*1^; *P <* .001). Note that the cascade of low *L*_*n*_ at the bottom to high negative *L*_*n*_ at the top of the canopy is equivalent to the cascade of thermal radiation exchange in the atmospheric layers and leads to the previously described ‘canopy greenhouse effect’ (Rotenberg and Yakir, 2011).

**Table 3.** Energy budget parameters (W m^*−*2^) at the bottom (*ca*. 2.5 m) and top of the canopy (*ca*. 10 m) for the projected leaf surface area during peak hours (11:00-14:00), where the heat storage (*G*) and fluorescence (*Fl*) were ignored due to their low magnitude (See Methods). *LE* at all heights was assumed to be the same as in the middle of the canopy, where it was measured. *H* was measured using sonic anemometers for layers of 1.5 m thickness, and was not significantly different between heights (*P* = 0.39) in spite of vast differences in short- and longwave radiation loads at the top and bottom of the canopy. Note that Sin is low even at the canopy top due to the lower halves of leaves that are shaded.

| Parameter | Bottom | Top |
| --- | --- | --- |
| $S_{in}$ | 345±142 | 488±69 |
| $S_{abs}$ | 241±99 | 341±48 |
| $L_{in}$ | 512±16 | 432±8 |
| $L_{abs}$ | 485±15 | 409±8 |
| $L_{emitted}$ | 473±7 | 465±10 |
| $L_n$ | 12±11 | -57±6 |
| $R_n$ | 253±109 | 285±52 |
| $LE$ | 3±3 | 8±13 |
| $H$ | 250±112 | 277±49 |

Lastly, leaves at the top were cooled convectively (*H*) while profiting from high Sin and low Lin, while leaves at the bottom of the canopy were exposed to a lower level of shortwave radiation and higher air temperature (by *ca*. 200 W m^*−*2^ and 2 °C, cf. Muller et al., 2021a). This reduces the ability of the latter to perform photosynthesis, even when soil water conditions improve. With a relatively stable Δ*T*_*L−A*_, *r*_*H*_ would also be affected, independently of the wind speed profile in the canopy. This level of variation within the canopy shows that the energy budget of a leaf in just one part of the canopy is insufficient to accurately represent heat exchange processes in the entire canopy. Nevertheless, this canopy-wide assessment of the twig-scale leaf energy budget is a fundamental tool to help improve modelling of ecosystem activity, specifically heat and gas exchange with the atmosphere and its effect on the climate system.

### Future improvements

As in-situ energy budget analyses, such as reported here, are still rare, we can point out to a number of caveats that can help improve future attempts at obtaining in-situ leaf-scale energy budgets: (a) Reflectivity and transmissivity were not measured for our needle-leaves, as they are difficult to obtain due to their small diameter. Development of appropriate methods will reduce uncertainties. (b) Neither our values of the incoming and outgoing *S* nor of *L* were based on the full spherical measurements, and this could be improved by adding a set of small hemispherical sensors near the leaves. In this study this was avoided as no sensors could be obtained that are small enough not to disturb the leaves. (c) The spectral attenuation of canopy interception could be measured using spectrometers near the leaves. This complicated and expensive setup was not available for our study, but could greatly improve future studies. (d) As *H* can’t currently be measured directly in the field, we propose a set of miniature air temperature thermocouples that aren’t affected by solar radiation, arranged in a 3D grid in and outside the leaf boundary layer, along with near-leaf wind measurements. This could help to fully assess sensible heat transport from the leaf to the mixed air in future studies.

## Supporting information

Full Supplement

## Acknowledgements

The authors are grateful to Stanislaus J. Schymanski for providing valuable comments that helped improve the paper, as well as Amnon Cochavi, Huanhuan Wang, Efrat Schwartz, Irina Vishnevetsky and Mila Volkov for their help in various tasks.

This study was supported by JNF-KKL (10-10920-19), Minerva (#714147), the ISF (#1976/17), a research grant from the *Yotam project* and the *Weizmann Institute Sustainability and Energy Research Initiative* as well as a fellowship from the *Society of Swiss Friends of the Weizmann Institute*.

## Author contribution

The study was conceived by JM, ER and DY; JM carried out the measurements with the help of IO and FT; JM analysed the data under the guidance of ER and DY; JM, ER and DY contributed to the writing.

## Code and data availability

The data that support the findings of this study are available from the corresponding author upon reasonable request. The software script developed for processing gas exchange chamber flux data is available as: ‘Branch-chamber-fluxes’ at https://doi.org/10.5281/zenodo.4284487, reference (Muller and Oz, 2020)

