## Supplementary material for "In-situ energy budget of needle-leaves reveals shift from evaporative to ‘air cooling’ under drought": Full Supplement

The following Supporting Information is available for this article:

**Methods S1:** Leaf surface area of needle-leaves

**Methods S2:** Latent heat & resistance to water transfer

**Methods S3:** Conversion of air molar volume to metric volume

**Methods S4:** Heat storage flux

**Methods S5:** Heat flux from sap flow into leaves ($F_S$)

**Methods S6:** Site description and meteorological conditions

**Methods S7:** Mean leaf thickness & transmittance

**Methods S8:** Workflow for absorbed shortwave radiation

**Methods S9:** Estimation of fluorescence

**Figure S1:** Comparison of *LOPEX93* and *ASD* measurements

**Figure S2:** Diurnal pattern of assimilation

**Notes S1:** Sensitivity of $H$ to air parameters

### Methods S1: Leaf surface area of needle-leaves

Needle-leaves of *P. halepensis* are close to pairs of half-cylinders, whereby flat-leaf equivalent measurements of S and L fluxes have to be adjusted to the total leaf surface area. The total leaf surface area ($LA$) of a single needle-leaf can be calculated from measurement of needle-leaf length ($l$) and diameter ($d$), where a fascicle (bundle) of needle-leaves forms a cylinder, and $n$ is the number of needles per fascicle (Grace, 1987), i.e. 2 for *P. halepensis*:

$$LA = \frac{\pi dl + ndl}{n} \tag{M.S1.1}$$

Note that measurements of incident radiation are typically per $m^2$ of flat surface area, i.e. equivalent to flat leaves. To adjust flat-leaf equivalent measurements to the total leaf surface area of half-cylinder leaves, flat-leaf equivalent measurements need to be corrected by a factor $x$, following:

$$2dlx = \frac{\pi dl + 2dl}{2} \tag{M.S1.2}$$

This results in a factor $x$ of:

$$x = \frac{\pi + 2}{4} \approx 1.3 \tag{M.S1.3}$$

### Methods S2: Latent heat & resistance to water transfer

Latent heat corresponds to the difference in water concentration between the leaf ($C_{w,L}$; $\mathrm{mol_{H_2O}\,m_{air}^{-3}}$) and the ambient air ($C_{w,A}$) divided by the resistance to water vapour transfer ($r_w$;$\mathrm{s\,m^{-1}}$):

$$LE = \lambda_w \, \frac{C_{w,L} - C_{w,A}}{r_w} \tag{M.S2.1}$$

A polynomial curve fit of measured values (Yau and Rogers, 1996; Qu et al., 2016), valid from −25 to 40 °C, yields an empirical equation that describes the dependence of the enthalpy of vaporisation ($\lambda_w$) on leaf temperature ($T_L$; °C), and the molar weight of water ($M_w$; 18.015 28 $\mathrm{g\,mol^{-1}}$):

$$\lambda_w = (2500.8 - 2.36\ T_L + 0.0016\ T_L^2 - 0.00006\ T_L^3)M_w \tag{M.S2.2}$$

Thus, the total resistance to water transfer ($r_w$) can be obtained from Eq. M.S2.1 using the evaporative flux ($E$, $\mathrm{mol_{H_2O}\,mol_{air}^{-1}}$) and the difference in water concentrations between leaf and air. $r_w$ consists of both the stomatal and the boundary layer resistance. However, the same equation for $r_w$ (below) yields the boundary layer resistance if an air water concentration near leaves is used for $C_{w,L}$ (e.g. inside branch chambers) instead of that inside the leaf.

$$r_w = \frac{C_{w,L} - C_{w,A}}{E} \tag{M.S2.3}$$

Details on obtaining $C_{w,L}$ and $C_{w,A}$ from the typically measured molar volumes are given in Methods S3.

### Methods S3: Conversion of air molar volume to metric volume

Water concentration in the air is measured by infrared gas analysers in mole fractions ($w_e$; $\text{mol}_{\text{H}_2\text{O}}\,\text{mol}_{\text{air}}^{-1}$), but is required as $C_{w,A}$ in $\text{mol}_{\text{H}_2\text{O}}\,\text{m}_{\text{air}}^{-3}$. Conversion can be achieved by dividing $w_e$ with the total molar volume of air ($v_a$; $\text{m}_{\text{air}}^{3}\,\text{mol}_{\text{air}}^{-1}$):

$$C_{w,A} = \frac{w_e}{v_a} \tag{M.S3.1}$$

$v_a$ is a function of the molar volume of dry air ($v_d$; $\text{m}_{\text{air}}^{3}\,\text{mol}_{\text{air}}^{-1}$) and the partial pressure of dry air ($P_d$; Pa) over the total air pressure ($P_a$; Pa):

$$v_a = v_d \frac{P_d}{P_a} \tag{M.S3.2}$$

$v_d$ can be obtained from the molecular weight of dry air ($M_d = 0.02897$ $\text{kg}\,\text{mol}^{-1}$) and its dry density ($\rho_d$; $\text{kg}\,\text{m}^{-3}$):

$$v_d = \frac{M_d}{\rho_d} \tag{M.S3.3}$$

$\rho_d$ is obtained from $P_d$, $M_d$, air temperature ($T_{air}$; K) and the ideal gas constant ($R = 8.314463$ $\text{J}\,\text{K}^{-1}\,\text{mol}^{-1}$):

$$\rho_d = \frac{P_d\, M_d}{R\, T_A} \tag{M.S3.4}$$

$P_d$ is the dry air partial pressure, obtained from the saturation vapour pressure ($e_s$; Pa):

$$P_d = P_a - e_s \tag{M.S3.5}$$

The saturation vapour pressure ($e_s$) has been described by Campbell and Norman (2012) using an empirical equation based on air temperature:

$$e_s = T_A^{-8.2} exp(77.345 + 0.0057\, T_A - 7235\, T_A^{-1}) \tag{M.S3.6}$$

$C_{w,L}$ is calculated for saturated air, which is assumed to be the condition inside the leaf:

$$C_{w,L} = \frac{w_{e,sat}}{v_{a,sat}} \tag{M.S3.7}$$

The saturated mole fraction ($w_{e,sat}$; $\mathrm{mol_{H_2O}\,mol_{air}^{-1}}$) has to be calculated for saturation ($RH = 100$ %), from the density of saturated air ($\rho_{A,sat}$; $\mathrm{kg\,m^{-3}}$) and the molecular weight of water vapour ($M_w = 0.01802\ \mathrm{kg\,mol^{-1}}$):

$$w_{e,sat} = \frac{\rho_{A,sat}\ v_{a,sat}}{M_w} \tag{M.S3.8}$$

$\rho_{A,sat}$ corresponds to the vapour pressure, here for saturated air, divided by the gas constant for water vapour ($R_w$; $\mathrm{J\,K^{-1}\,kg^{-1}}$) and leaf temperature ($T_L$; K):

$$\rho_{A,sat} = \frac{e_s}{R_w\ T_L} \tag{M.S3.9}$$

$R_w$ can be obtained from the universal gas constant ($R$; $\mathrm{J\,K^{-1}\,mol^{-1}}$):

$$R_w = \frac{R}{M_w} \tag{M.S3.10}$$

Finally, plugging all the above into Eq. M.S3.7 yields:

$$C_{w,L} = \frac{e_s}{R\ T_L} \tag{M.S3.11}$$

### Methods S4: Heat storage flux

$G$ is a function of the change in $T_L$ over time (i.e., $dT_L/dt$) resulting from heat transferred per total leaf surface area (LA) into the mass of a leaf, i.e., that of water ($m_w$) and carbon ($m_C$) contained in it. The specific heat capacity of water ($c_{p,w}$) and of carbon ($c_{p,C}$) at 20 °C is 4.182 and 0.71 $\mathrm{J\,K^{-1}\,g^{-1}}$, respectively:

$$G = \frac{c_{p,w}\ m_w + c_{p,C}\ m_C}{LA}\ \frac{dT_L}{dt} \tag{M.S4.1}$$

Note that some studies simply assume that the overall leaf heat capacity, $c_{p,L}$, can be sufficiently accurately simplified by $c_{p,w}$, considering that leaf water content is usually 80 to 90 % and has a high heat capacity (e.g., Schymanski et al., 2013). Finally, $G$ is negligibly small in small leaves, but it can be considerable in thick leaves.

Needle-leaf wet and dry weight were measured monthly in irrigated and drought-exposed plots, both in pre-dawn and midday conditions. Due to the small dimensions of needle-leaves (here 0.8 to 1 mm diameter), the leaf temperature was assumed to be equivalent to the leaf surface temperature. The resulting $G$ per leaf surface area (Eq. M.S4.1) was negligible (mean of $0.06 \pm 0.16\ \mathrm{W\,m^{-2}}$, but note that for short-term sunspots appearing and disappearing in at short time scales, $G$ could be a large component.

The ratio of $G$ to $LE$ for a high transpiration rates of $0.5\,\mathrm{mmol\,m^{-2}\,s^{-1}}$, as are found in irrigated trees at the Yatir research site, was less than 0.01, showing that this is a minor contribution to the leaf energy budget.

### Methods S5: Heat flux from sap flow into leaves ($F_S$)

As water evaporates from leaves, it is replenished with sap coming from branches. This sap may have a different temperature ($T_S$) than leaves ($T_L$): Branches heat up considerably when exposed to sunlight, but their bark also insulates the sap from this heating effect. Assuming that the flow rate of sap into a leaf is equivalent to the transpiration rate (TR; $\mathrm{mol\,m^{-2}\,s^{-1}}$), and that the heat capacity and molar weight of sap are equivalent to those of water (i.e., $c_{p,w}$ and $M_w$), the heat flux from leaves gained or lost through this sap flow ($F_S$) can be formulated as follows:

$$F_S = c_{p,w}\ M_w(T_L - T_S)\ TR \tag{M.S5.1}$$

Since branch sap temperature was not available in the current study, a sensitivity analysis is provided in Fig. M.S5.1, showing the sensitivity of energy lost or gained due to sap and leaf temperature differences from −5 to 5 °C, for transpiration rates ranging from 0 to $0.5\,\mathrm{mmol\,m^{-2}\,s^{-1}}$: Results show that during high transpiration rates of $0.5\,\mathrm{mmol\,m^{-2}\,s^{-1}}$, as are found in irrigated trees at the Yatir research site, and a sap temperature *ca.*2.5 °C higher than leaf temperature, around $0.1\,\mathrm{W\,m^{-2}}$ is gained by the leaf. In our conditions, branch/twig surface temperature was often this much warmer than leaves, but as the bark acts as an insulator, the exact sap temperature was unknown.

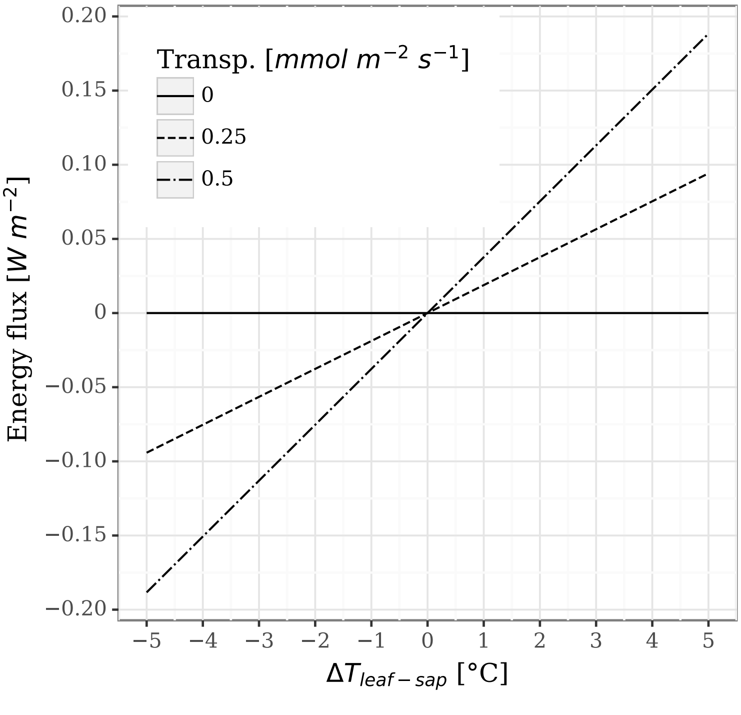


**Fig. M.S5.1.** Sensitivity of energy lost or gained due to sap and leaf temperature differences from −5 to 5 °C, for transpiration rates ranging from 0 to $0.5\,\mathrm{mmol\,m^{-2}\,s^{-1}}$

### Methods S6: Site description and meteorological conditions during measurement periods

The Yatir forest research site is located in the dry southern Mediterranean region, at the northern edge of the Negev desert in Israel (31°20′49″N; 35°3′7″E; altitude 600 to 850 m above sea level) Tree density is 300 trees $ha^{-1}$ (Tatarinov et al., 2016), and the lowest branches were at ~2.5 m agl. Mean annual global radiation was 238 $W\,m^{-2}$ (Rotenberg and Yakir, 2010), while average air temperatures for January and July are 10 and 25.8 °C respectively, with a mean annual potential ET of 1600 mm, and mean annual precipitation of 285 mm (Qubaja et al., 2019). This ecosystem is exposed to an extended dry season (May to October) typical for the semi-arid southern Mediterranean climate zone (Qubaja et al., 2020).

Mean and standard deviations of midday conditions (10:00-14:00; Table M.S6.1) were similar during the consecutive measurement periods: air temperature was *ca.* 30 °C, above-canopy incoming short- and longwave radiation were *ca.* 900 and 360 $W\,m^{-2}$, respectively, while above-canopy (4 m above canopy) wind speed averaged 3.2 $m\,s^{-1}$. These values show that conditions during summer drought remain stable and comparable, even though measurements were not taken simultaneously in the two plots. Needle leaf diameter was *ca.* 0.8 to 1 mm, while length was *ca.* 70 mm in drought-exposed and *ca.* 85 mm in irrigated trees at 5 m agl (supplement of Muller et al., 2021).

**Table M.S6.1.** Mean and standard deviations of midday conditions (10:00-14:00) during consecutive measurements periods in summer 2019 with standard deviations, showing drought-exposed ecosystem-scale meteorological conditions measured above canopy at 15 m agl for the two consecutive measurement periods in different treatments (irrigated and drought-exposed). Columns show the exact dates of measurements, air temperature $T_{air}$ (°C), down- (↓) and up-welling (↑) shortwave ($S$) and longwave thermal radiation ($L$; $W\,m^{-2}$), sensible ($H$; $W\,m^{-2}$) and latent heat flux ($LE$; $W\,m^{-2}$) and wind speed $u$ ($m\,s^{-1}$), respectively.

| Treatment<br>Dates | Drought-exposed<br>25.6. − 31.7. | Irrigated<br>1.8. − 30.9. |
|---|---|---|
| $T_A$ | 29.58±2.24 | 30.18±2.15 |
| $S_{\downarrow,ac}$ | 842±130 | 935±96 |
| $L_{\downarrow,ac}$ | 362±16 | 359±14 |
| $S_{\uparrow,ac}$ | 106±14 | 114±11 |
| $L_{\uparrow,ac}$ | 500±16 | 508±14 |
| $H$ | 531±84 | 493±85 |
| $LE$ | 36±38 | 36±38 |
| $u$ | 2.97±0.84 | 3.26±0.88 |

### Methods S7: Mean leaf thickness & transmittance

Radiation load on leaves depends on absorptance, which in itself is the remaining fraction after reflectance and transmittance. Broadleaves reduce absorptance by being thinner and more transmissive, while needle-leaves are thicker but thinner, i.e. have less surface area that absorbs that radiation. Additionally, leaf angle, shape, structure, chlorophyll content and other factors affect both reflectance and absorptance. However, these parameters could not be taken into account in this study.

Transmittance is inversely correlated to leaf thickness, i.e. the thicker a leaf, the less radiation is transmitted. To account for needle-leaf transmittance, the transmittance $\tau_{s,\lambda}$ at each wavelength of each fresh leaf in the LOPEX93 database (Hosgood et al., 1995) was normalised using its thickness $T$ to the mean width of a needle-leaf.

$$\tau_{s,\lambda,norm} = \frac{\tau_{s,\lambda}\ T}{W} \qquad \text{(M.S7.1)}$$

Then, the normalised transmittance $\tau_{s,\lambda,norm}$ was averaging across all species for each wavelength of the spectrum.

The mean thickness $W$ of a needle-leaf was approximated by the mean width of a circle: $W$ is the area of the circle divided its diameter.

$$W = \frac{\pi\ r^2}{2\ r} = \frac{1}{2}\pi\ r \qquad \text{(M.S7.2)}$$

For our near-circular *P. halepensis* needle-leaves of 0.8 mm diameter, this corresponded to a mean thickness of ca. *ca.* 0.62 mm.

### Methods S8: Workflow for absorbed shortwave radiation

The absorbed radiation at the leaf ($S_{abs}$) was calculated according to the workflow presented in Fig. M.S8.1 from the radiation absorbed by the upper and lower halves of a leaf ($S_{abs,upper}$ and $S_{abs,lower}$).

**Downwelling spectral radiant flux at the leaf:** $S_{\lambda,\downarrow,aL}$ corresponds to the above-canopy downwelling $S_{\lambda,\downarrow,ac}$, adjusted to measurements of $S_{\downarrow,aL}$ from the photodiode to take the canopy attenuation into account. Note that theoretically, $S_{\lambda,\downarrow,aL}$ is not identical to $S_{\lambda,\downarrow,ac}$ since the canopy modifies the spectrum and doesn't just attenuate it. However, this could not be taken into account here:

$$S_{\lambda,\downarrow,aL} = S_{\lambda,\downarrow,ac}\ \frac{S_{\downarrow,aL}}{\sum S_{\lambda,\downarrow,ac}} \qquad \text{(M.S8.1)}$$

$S_{\lambda,\downarrow,ac}$ was obtained from a calibration of the modelled $S_{\lambda,\downarrow,mod}$, adjusted to the measurements of $S_{\downarrow,ac}$ (Eq. M.S8.2) in order to account for model inaccuracies from clouds and airborne particles such as dust. $S_{\lambda,\downarrow,mod}$ was estimated using the *SMARTS2* clear-sky spectral model (Gueymard, 2005, 1995; Myers and Gueymard, 2004). *SMARTS2* was initialised with environmental data measured above canopy at our Eddy Covariance

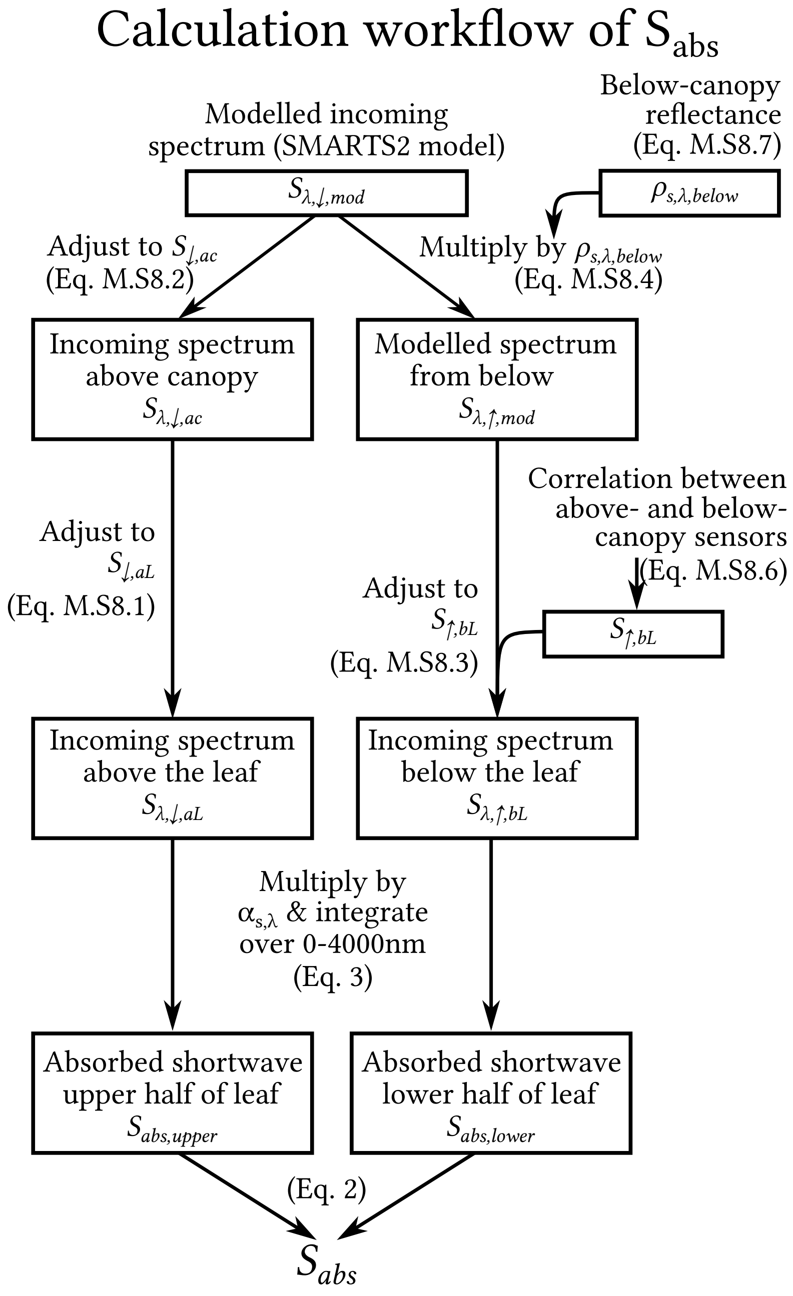


**Fig. M.S8.1.** Flowchart of calculation of shortwave radiation absorbed by a leaf ($S_{abs}$) from measured above- ($S_{\downarrow,ac}$, $S_{\uparrow,ac}$), below- ($S_{\downarrow,bc}$, $S_{\uparrow,bc}$ and $\rho_{s,\lambda,below}$) and within-canopy variables ($S_{\downarrow,aL}$), literature values of the leaf absorptance spectrum ($\alpha_{s,\lambda}$), and the modelled incoming light spectrum above the canopy ($S_{\lambda,\downarrow,mod}$)

flux tower, i.e. latitude, longitude, altitude, date & time, air temperature, relative humidity, atmospheric $CO_2$ concentration and ecosystem albedo.

$$S_{\lambda,\downarrow,ac} = S_{\lambda,\downarrow,mod} \frac{S_{\downarrow,ac}}{\sum S_{\lambda,\downarrow,mod}} \tag{M.S8.2}$$

The $S_{\downarrow,ac}$ used here were interpolated to 6min time intervals from the available half-hourly data measured at the Eddy Covariance tower. This 6min time interval was identified as the appropriate averaging period for turbulence and wind measurements within the canopy in our conditions and at other sites (Thomas et al., 2013). Note that for diurnal graphs, all data at the 6min resolution was averaged to half-hours to reduce the noise.

**Upwelling shortwave spectral radiant flux at the leaf:** $S_{\lambda,\uparrow,bL}$ was estimated from the modelled $S_{\lambda,\uparrow,mod}$, calibrated by $S_{\uparrow,bL}$ (Eq. M.S8.3).

$$S_{\lambda,\uparrow,bL} = S_{\lambda,\uparrow,mod} \frac{S_{\uparrow,bL}}{\sum S_{\lambda,\uparrow,mod}} \quad \text{(M.S8.3)}$$

$S_{\lambda,\uparrow,mod}$ corresponds to the modelled downwelling $S_{\lambda,\downarrow,mod}$ from the sky, modified by the reflectance spectrum of the canopy and ground from which it is reflected from below the measured leaf ($\rho_{s,\lambda,below}$; Eq. M.S8.4).

$$S_{\lambda,\uparrow,mod} = S_{\lambda,\downarrow,mod}\ \rho_{s,\lambda,below} \quad \text{(M.S8.4)}$$

*Estimation of total upwelling shortwave radiation in the canopy:* $S_{\uparrow,bL}$ was estimated empirically using a correlation derived from measurements at the Eddy Covariance tower (*ca.* 50 m away from energy budget measurements). The downwelling $S_{\downarrow}$ (mean of above and below canopy: $S_{\uparrow,ac}$ and $S_{\uparrow,bc}$) was correlated with the reflected upwelling $S_{\uparrow}$ (mean of above and below: $S_{\downarrow,ac}$ and $S_{\downarrow,bc}$; Eq. M.S8.5) through a linear equation with a slope $d$ and an intercept $e$ ($d = 0.16$ and $e = 0.54$, $R^2 = 0.98$, $P < .001$; $RMSE = 4.90$ $\mathrm{W\,m^{-2}}$), as shown in Fig. M.S8.2. This correlation was used to estimate upwelling S at the location of leaf energy budget measurements.

$$\frac{S_{\downarrow,ac} + S_{\downarrow,bc}}{2} = d\ \frac{S_{\uparrow,ac} + S_{\uparrow,bc}}{2} + e \quad \text{(M.S8.5)}$$

Eq. M.S8.5 was then solved for the upwelling component at the measurement location ($S_{\uparrow,bc}$, which corresponds to the required $S_{\uparrow,bL}$). The above-canopy $S_{\downarrow,ac}$ and $S_{\uparrow,ac}$ were retained while the downwelling component below canopy ($S_{\downarrow,bc}$) was replaced with data from the measurement location (i.e., $S_{\downarrow,aL}$):

$$S_{\uparrow,bL} = \frac{S_{\downarrow,ac} + S_{\downarrow,aL}}{d} - 2\ \frac{e}{d} - S_{\uparrow,bc} \quad \text{(M.S8.6)}$$

Note that due to the small magnitude of $S_{\uparrow,bL}$, inaccuracies in its estimation will have a minor effect on overall results.

*Estimation of within-canopy reflectance:* $\rho_{s,\lambda,below}$ was estimated to be the mean spectral reflectance of the canopy ($\rho_{s,\lambda,canopy}$) and ground ($\rho_{s,\lambda,ground}$), and each of those is composed of a combination of the reflectance of different materials (young and old leaves, soil, stones and plant litter) in proportions approximately estimated for the Yatir forest research site. Reflectance spectra for each material were measured using an ASD spectrometer (ASD FieldSpec 4, Malvern Panalytical; Malvern, United Kingdom). Note that the weighted mean of the $\rho_{s,\lambda,ground}$ spectrum was *ca.* 0.35. However, a rough estimation using daytime below-canopy up- and downwelling measurements (i.e. $\rho_{ground,est} = S_{\uparrow,bc}/S_{\downarrow,bc}$) yielded a value of $0.17 \pm 0.06$ across an entire month. Therefore, the $\rho_{s,\lambda,ground}$ spectrum was adjusted by a factor 0.5. ASD measurements of the needle

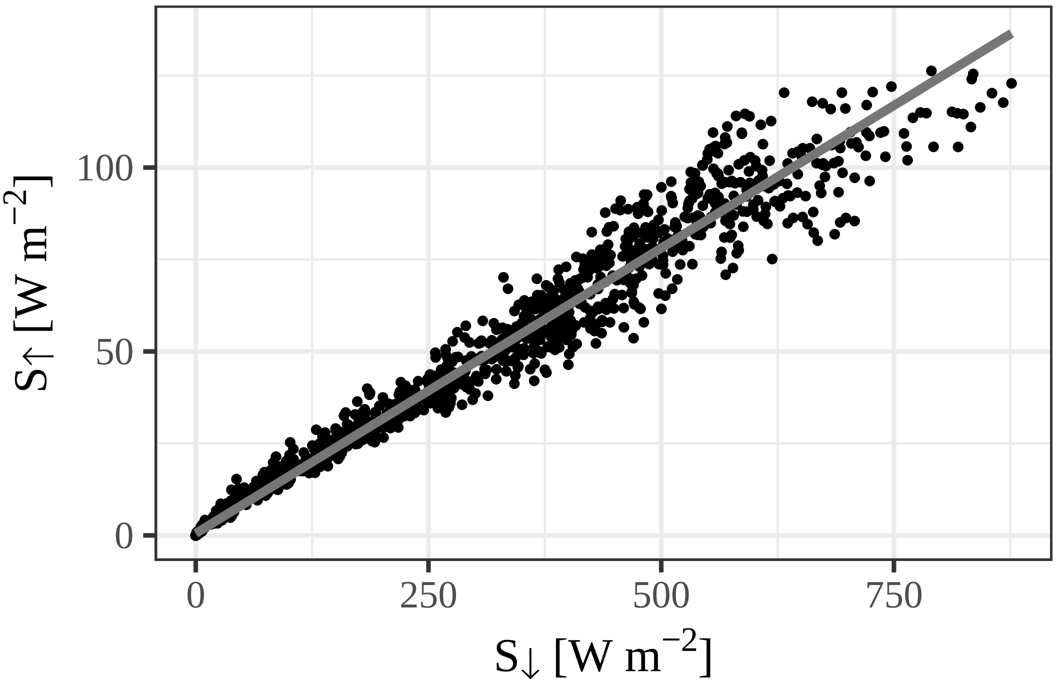


**Fig. M.S8.2.** Correlation between downwelling $S_\downarrow$ (mean of above and below canopy) and reflected upwelling $S_\uparrow$ (mean of above and below), with $S_\uparrow = 0.16\ S_\downarrow + 0.54$; $R^2 = 0.98$, $P < .001$; $RMSE = 4.90\ \mathrm{W\,m^{-2}}$

reflectance spectra were used here. Note that were within the standard deviation range of the *LOPEX93* database values for the most energetic part of the spectrum (350 to 1400 nm; Figure S3).

$$\begin{aligned} \rho_{s,\lambda,canopy} &= 0.5\ \rho_{s,\lambda,young\ needles} + 0.5\ \rho_{s,\lambda,old\ needles} \\ \rho_{s,\lambda,ground} &= 0.5\ (0.25\ \rho_{s,\lambda,soil} + 0.15\ \rho_{s,\lambda,stones} + 0.65\ \rho_{s,\lambda,litter}) \end{aligned} \quad \text{(M.S8.7)}$$

### Methods S9: Estimation of fluorescence

Amount of energy emitted through fluorescence estimated using a simulation with the Soil-Canopy-Observation of Photosynthesis and Energy fluxes (SCOPE) model (v.1.7; van der Tol et al., 2009; https://github.com/Christiaanvandertol/SCOPE), initialised with data from the Eddy Covariance tower. This model is widely used and has previously been shown as appropriate for pine forests (Rossini et al., 2016). Results are shown in Fig. M.S9.1.

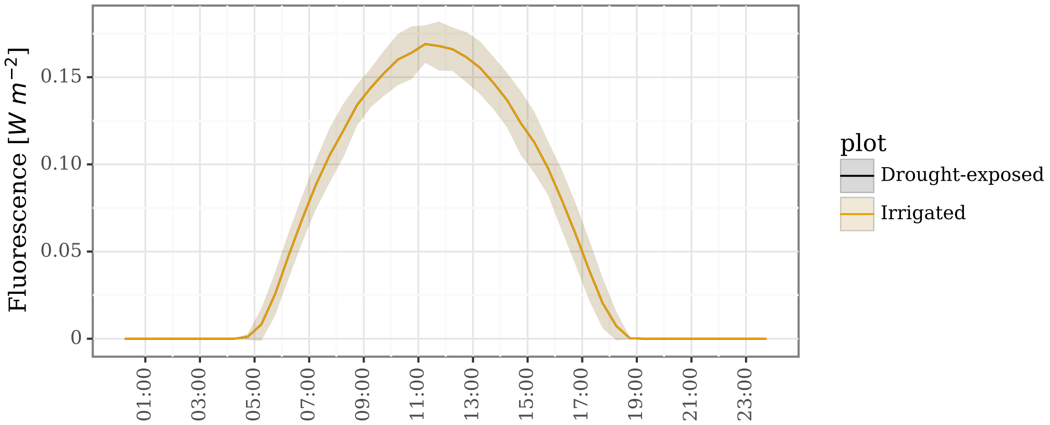


**Fig. M.S9.1.** Simulation of fluorescence as a diurnal trend in drought-exposed and irrigated treatments (Note: The curve of the drought-exposed treatment is under the one of the irrigated one). Simulations were done using the SCOPE model, where shading represents standard deviations.

### Figure S1: Comparison of *LOPEX93* and *ASD* measurements

Reflectance mean of all broadleaf species in the *LOPEX93* database (Hosgood et al., 1995) vs. ASD reflectance spectrum measurements (ASD FieldSpec 4, Malvern Panalytical; Malvern, United Kingdom) of young and old needles, showing subtle differences between both measurements of reflectance. The ASD measurements were not hemispherical and only provided reflectance, not transmittance. Therefore, ASD measurements were only used as a reference to qualitatively check whether *LOPEX93* values were reasonable, showing that they were indeed comparable. Note: While reflectance data is available for needle-leaves in the LOPEX93 database, transmittance data is not. Therefore, the mean of all broadleaf species was used in the current study, and needle-leaves were omitted from the figure as a means of comparison to the rest of the study.

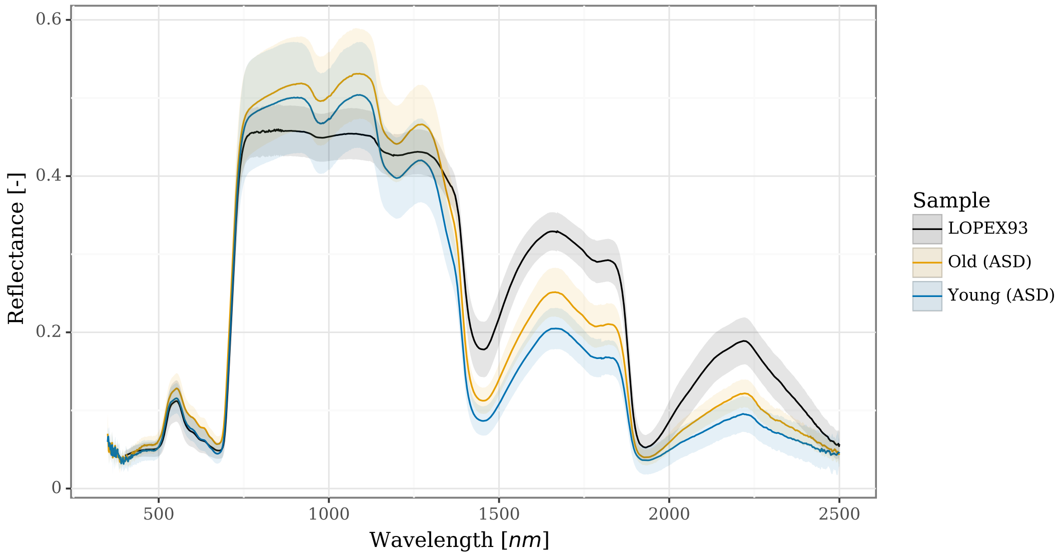

### Figure S2: Diurnal pattern of assimilation

The diurnal pattern of assimilation, adapted from Preisler (2019) and Oz (2021), is shown in 30 min time steps for the droughted (black) and irrigated treatments (yellow), with shaded areas representing standard deviations. The flux is shown in the direction from the leaf to the atmosphere, i.e. negative values represent a $CO_2$ uptake. Assimilation remained near zero in the droughted treatment, except during early morning hours, while it was up to 7× larger in irrigated trees.

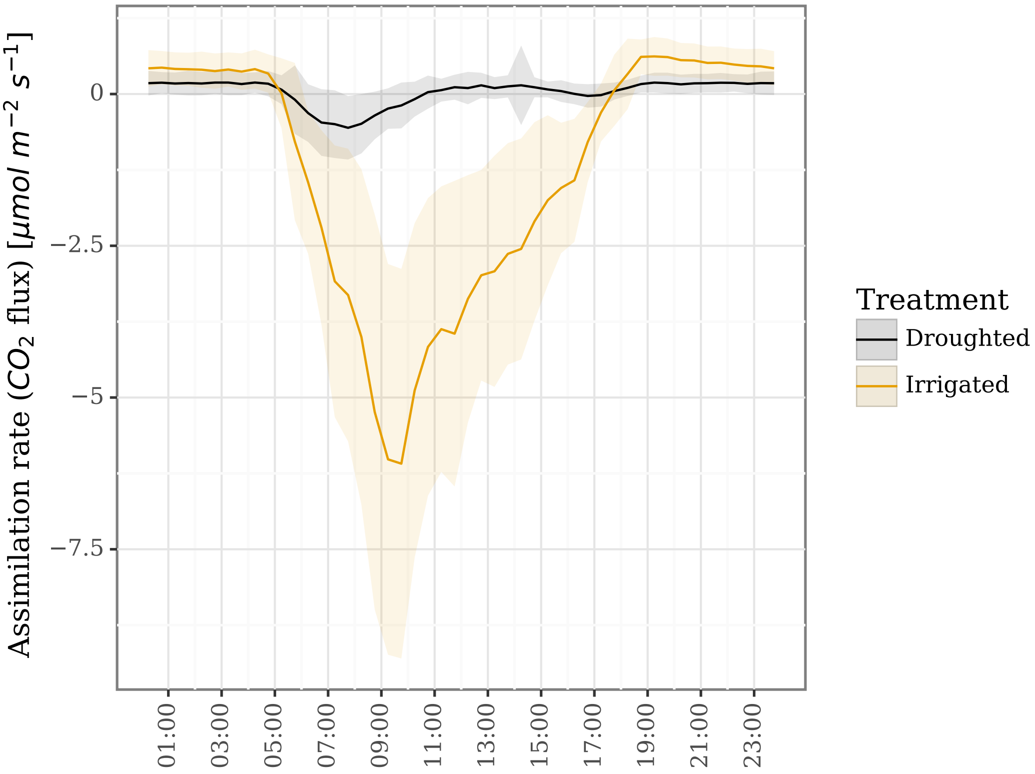


### Notes S1: Sensitivity of $H$ to air parameters

The moisture content of the air surrounding leaves, resulting from transpiration, raise of lower $H$ through its effect on the heat capacity $c_{p,A}$ and density $\rho_A$ of the air. A sensitivity analysis was done where the following parameters were kept constant in the sensible heat equation (Eq. 5): $T_A$ = 30°C, $P_A$ = 934 hPa, $\Delta T_{L-A}$ = 3°C, $r_H$ = 20 $\mathrm{s\,m^{-1}}$. Then, air moisture content was changed from 20 to 35 $\mathrm{mmol\,mol^{-1}}$ to affect $c_{p,A}$ and $\rho_A$, i.e. increasing them by 8.5 $\mathrm{J\,kg^{-1}\,K^{-1}}$ and 0.01 $\mathrm{kg\,m^{-3}}$, respectively. This resulted in an $H$ difference of merely *ca.* 2.8 $\mathrm{W\,m^{-2}}$.
